# Predictive *in vitro* profiling of LNP-induced innate immune response using an iPSC-derived monocyte model

**DOI:** 10.1101/2025.09.26.678842

**Authors:** Baixue Xiao, Soojin Kim, Benjamin I Laufer, Anthony Antonelli, Milena Hornburg, Divya Murali, Yimin Gu, Pierce Jessen, Diamanda Rigas, Rebecca Leylek, Keiko Hokeness, Emily Freund, Eric Torres, Gaia Ruggeri, Siri Tahtinen, Yuchen Fan, Chun-Wan Yen, Maheswara Reddy Emani

**Affiliations:** Synthetic Molecule Pharmaceutical Sciences, Genentech, South San Francisco, CA 94080, USA; Biochemical and Cellular Pharmacology, Genentech, South San Francisco, CA 94080, USA; Computational Biology & Medicine, Genentech, South San Francisco, CA 94080, USA; Molecular Oncology, Genentech, South San Francisco, CA 94080, USA; Immunology Discovery, Genentech, South San Francisco, CA 94080, USA; Molecular Biology, Genentech, South San Francisco, CA 94080, USA

## Abstract

Lipid nanoparticles (LNPs) are a powerful drug delivery platform advancing vaccines and gene therapies. While their efficacy and safety has been found to be closely linked to innate immune activation, current in vitro models are unable to predict immune responses reliably. Conventional models, such as PBMCs, are limited by donor variability and inconsistent sensitivity. To address this, we developed a cytokine profiling platform using induced pluripotent stem cell (iPSC)-derived monocytes (iMonocytes), a physiologically relevant innate immune cell type that plays a key role in immune surveillance and inflammation. iPSCs provide a renewable, uniform monocyte source for consistent, high-sensitivity LNP screening. When tested with LNPs of graded immunostimulatory potency, iMonocytes showed improved reproducibility and strong correlation with *in vivo* cytokine responses. This platform enables evaluation of cargo- and dose-dependent effects, providing a robust and scalable tool for preclinical assessment and rational design of LNP therapeutics.

## Introduction

Lipid nanoparticles (LNPs) have revolutionized the landscape of modern medicine. This versatile drug delivery platform has enabled breakthroughs in vaccines, cancer immunotherapies, and gene therapies due to its rapid design, scalable manufacturing, and ability to encapsulate diverse therapeutic cargos, particularly nucleic acids, which are traditionally challenging to be safe and effectively delivered.^1–5^ A critical determinant of the success of LNP platforms lies in their interaction with the innate immune system, a double-edged sword that presents both opportunities and challenges.^6–12^ For sustained efficacy of protein replacement or gene editing therapies targeting genetic disorders like cystic fibrosis, minimizing innate immune recognition is crucial to prevent unnecessary inflammation and neutralizing antibody formation.^13^ Conversely, vaccine development requires precisely tuned innate immune activation to elicit robust adaptive responses;^14, 15^ while not triggering excessive inflammation to compromise safety and/or effectiveness. Therefore, as the LNP field rapidly advances, it becomes increasingly important to understand and control potential innate immune responses triggered by LNPs to ensure both efficacy and safety across this expanding medical landscape.

The innate immunostimulatory potential of LNPs exhibits substantial variability depending on LNP composition, including the ionizable lipids, structural lipids, PEGylation, steroid lipids, and overall physicochemical properties at the nanoparticle scale. Among these components, the synthetic, non-natural ionizable lipid serves as the primary determinant of immune modulation,^16^ demonstrated the critical role of ionizable lipids in mediating LNPs’ adjuvant effects, demonstrating their capacity to stimulate interleukin 6 (IL-6) production from innate immune cells and subsequently drive potent germinal center B-cell and T follicular helper cell responses. This mechanistic insight was expanded by Tahtinen et al, who demonstrated that lipid formulated nanoparticles, including LNPs, can induce cytokines such as IL-6 via inflammasome-mediated activation of interleukin 1 (IL-1),^17^ further studied LNP-induced innate immune responses by conducting a systematic comparison of six clinically relevant ionizable lipids (MC3, SM-102, ALC-0315, Moderna Lipid 5, Compound 9, and CKK-E12). Their work revealed striking differences in innate immune activation, with CKK-E12 emerging as the most potent inducer of monocyte chemoattractant protein-1 (MCP-1), eliciting significantly higher cytokine levels at six hours post-administration compared to other lipids.^17^ These findings underscore the importance of comprehensive lipid screening to enable rational design of LNP formulations with precisely tuned immunostimulatory properties. ^7, 8–12^

Standardized regulatory frameworks for assessing the immunostimulatory properties of LNP-based therapies remain lacking, necessitating the development of robust *in vitro* models for preclinical development. Primary human immune cells, such as peripheral blood mononuclear cells (PBMCs), including monocytes, are widely used to study LNP-induced innate immune responses due to their physiological relevance.^6^ However, inability to culture primary cells for prolonged time for kinetic studies and potential donor-to-donor variability compromises reproducibility;^18, 19^ while immortalized cell lines (e.g., THP-1), though scalable, exhibit altered morphology and limited stimulus responses.^20, 21^ To overcome these limitations, induced pluripotent stem cell (iPSC)-derived monocytes (iMonocytes) have emerged as a promising alternative. Human iPSCs offer an unlimited self-renewal and differentiation potential, enabling the generation of genetically uniform, functionally consistent myeloid cells for disease modeling, therapeutic development, and immunostimulatory response screening.^22–27^

Here, we present the development of an *in vitro* iMonocyte model to evaluate the innate immune responses of LNPs. We first evaluated the model performance for rank ordering several model LNPs formulated with ionizable lipids with potentially graded immunostimulatory potential: SM-102, MC3 and Lipid 20, eliciting low-to-intermediate innate immune responses; and CKK-E12, eliciting high innate immune responses. Using comprehensive cytokine profiling, a well-validated biomarker of innate immune activation, we demonstrated that iMonocytes show superior predictive value when benchmarked against conventional PBMC and primary monocyte assays. Importantly, our *in vitro* findings correlate strongly with *in vivo* cytokine data, validating the translational relevance of this platform. Beyond characterizing LNP vehicle associated cytokine release, we further showed the utility of iMonocytes for assessing cargo-dependent innate immunostimulatory properties and optimizing dosing regimens. iPSC-derived monocytes show great promise as a robust, scalable, and translational platform for screening and optimizing LNP-based therapeutics during preclinical development.

## Results

### iMonocytes recapitulate the phenotype and function of primary monocytes

We first assessed the phenotypic feature of iMonocytes. Characterization of two human iPSC lines, IP11 and SCTi003, revealed robust expression of pluripotency markers (NANOG, OCT4, SSEA4, and TRA-1-81) as confirmed by immunofluorescence staining (Figure S1a, b). Genomic integrity was further assessed via KaryoStat™ analysis, which demonstrated normal karyotypes and an absence of chromosomal aberrations in both lines (Figure S1c, d). Monocytes were derived from both human iPSC lines using an adapted differentiation protocol from STEMCELL Technologies.^28^ A schematic diagram illustrates the differentiation steps from iPSCs to functional monocytes, with representative cell morphologies at each stage (Figure. 1a). During the initial 7 days, hematopoietic stem cells (HSCs) were induced, characterized by the expression of CD34 (Figure S1e, f). From day 8 onwards, cells were maintained in monocyte-specific media and harvested for downstream assays between days 18 and 20. The harvested iMonocytes can be used at least a week, with consistent cell viability of ∼85% in this assay period (Figure 1b). Surface markers CD14 and CD16 were analyzed in the harvested cells, with ∼70% being CD14^+^CD16^+^ (Figure. 1c). In our optimized T175 flask differentiation format, we generated 25∼30 million CD14^+^ and CD16^+^ iMonocytes and produced 0.3-1.2 million cells per well in a 6-well plate, demonstrating both the scalability of iMonocyte production and the flexibility of our protocol to accommodate different production scales based on experimental needs. This protocol was validated using two independent human iPSC lines.

**Figure 1.**
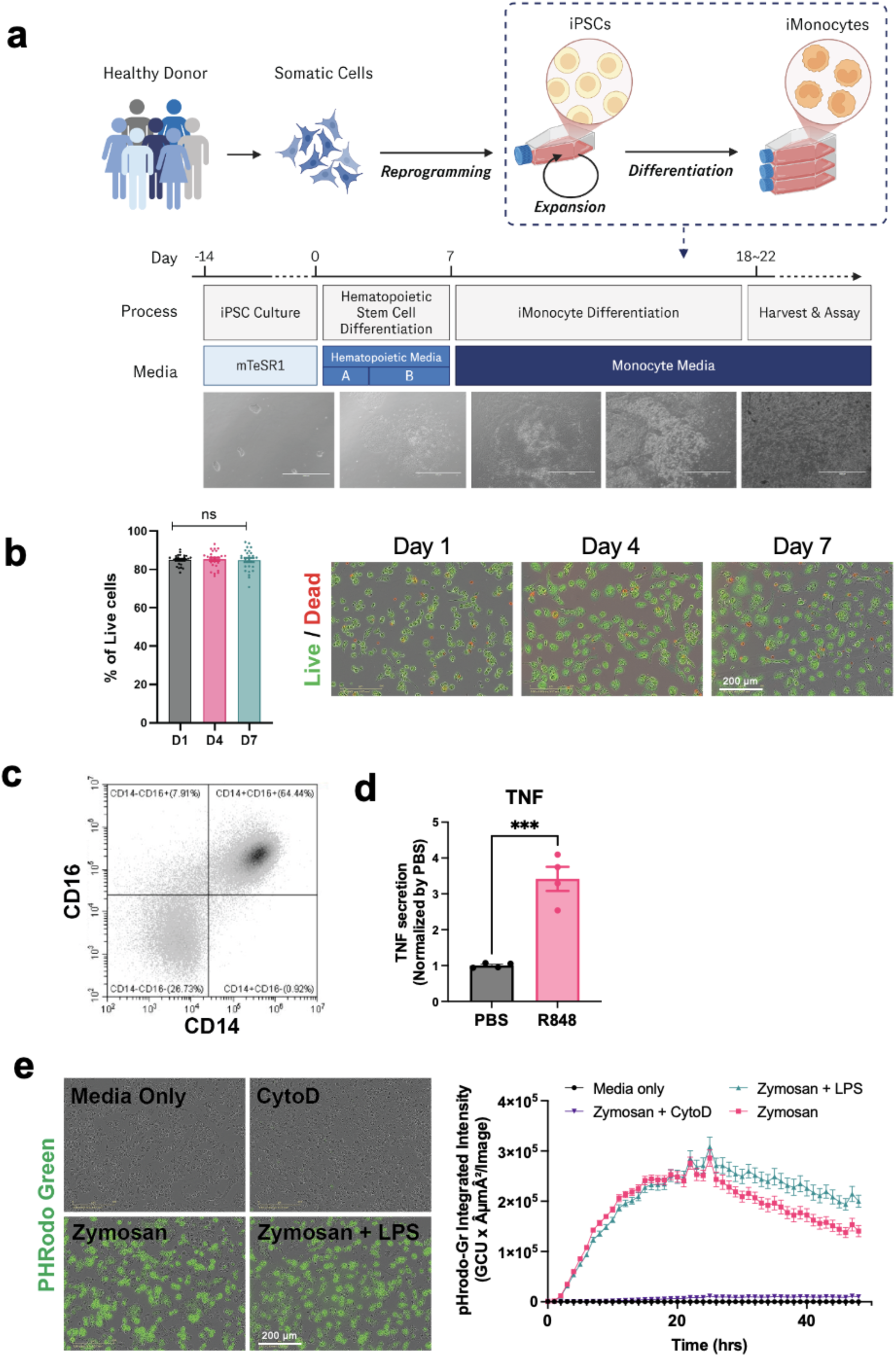
iPSC-derived monocytes recapitulate the phenotype and function of primary monocytes. **(a)** Schematic diagram of the differentiation procedure from iPSCs into functional monocytes. Cell morphology at each stage was also shown. **(b)** Live/dead cell viability assay of iMonocytes during the assay window at day 1, 3, and 7 post-harvest, showing consistent viability of ∼85% up to day 7. Statistic results were collected from 27 representative cell images at each time point. **(c)** Flow cytometry results showed ∼70% iPSC-derived monocytes were CD14 and CD16 double positive on day 7 post-harvest. **(d)** TNF-α release from iMonocytes following the R848 treatment (0.1 µg/mL for 16-18 hrs). Results were measured from n = 4, \*\*\**p* = 0.0004. **(e)** Phagocytosis of iMonocytes stimulated by zymosan-conjugated pHrodo Green bioparticles. Co-treatment of the phagocytosis inhibitor cytochalasinD (CytoD) or the inflammation inducer LPS were also tested. Representative cell images showed phagocytosis at 24 hrs post-stimulation (left). Total integrated intensity within cell boundaries was measured hourly for 48 hrs (right). Results show mean ± SEM, n = 4. Scale bars of cell images = 200 μm.

Upon the treatment of R848, which is a Toll Like Receptor (TLR) 7/8 agonist, iMonocyte secreted significant levels of TNF (Figure 1d). Further, iMonocyte also exhibited robust phagocytosis capability, as measured by the cellular uptake of pHrodo green-conjugated zymosan A particles (Figure 1e). The phagocytosis peaked at ∼24 hrs, completely inhibited by cytochalasin D (phagocytosis inhibitor by blocking the actin polymerization), while not significantly affected by the acute inflammation inducer, lipopolysaccharide (LPS). Overall, these results demonstrate iMonocytes recapitulate monocyte functionality, in terms of innate cytokine secretion and phagocytosis.

### R848 priming enhances the consistency and magnitude of mRNA-LNP-induced immune response in iMonocytes

To evaluate the iMonocytes as an *in vitro* platform for assessing the immune-stimulatory responses of LNPs, a panel of representative mRNA-LNP formulations was designed to span a spectrum of innate immune activation. Specifically, N1-methylpseudouridine (m1Ψ) modified mRNA-LNPs were prepared using typical composition ratios of the ionizable lipid, cholesterol, DOPE, and 14:0 PEG2000 PE. Four different ionizable lipids, MC3, SM102, Lipid 20, and CKK-E12, were selected to potentially provide distinct immunogenic profiles, ranging from SM102, MC3, Lipid 20 (low to moderate), to CKK-E12 (high) stimulation.^17^ SM102, MC3, and Lipid 20 mRNA-LNPs showed mean diameter ∼120 nm and encapsulation efficiency >95%; CKK-E12 mRNA-LNP showed slightly larger particle size (mean diameter of 138 nm) and slightly lower mRNA encapsulation (∼84%) (Figure S2a). All four types of mRNA-LNPs showed similar morphology and the structure feature of condensed cores, as measured by cryo-TEM (Figure S2b).

Interleukin 1 (IL-1) plays a critical role in regulating the innate immune response of monocytes to mRNA-LNP and may modulate the secretion of additional cytokines relevant to downstream analysis.^7, 10^ Building on our previous observation that detection of nanoparticle-induced inflammasome activation and IL-1 release is enhanced *in vitro* in the presence of signal 1 (TLR agonist such as R848,^7^ we sought to investigate if the sensitivity of our iMonocyte assay could be enhanced by pre-treating the cells with low-dose R848. iMonocytes were pre-treated with or without R848 for 2 hours, followed by washing out and LNP treatment for 3 days, and then measured for IL-1 secretion (Figure 2a, b) and mRNA translation (Figure S3). In the absence of R848, cytokine levels showed little difference between SM102 and Lipid 20, or between the LNPs and controls. Treatment with R848 alone led to a moderate, dose-independent IL-1α and IL-1β responses. The R848 dose titration results indicated that pre-treatment with a narrow dose window around 0.1 μg/mL allowed clear differentiation in IL-1 cytokine responses induced by different LNPs (Figure 2a, b). Further increasing the R848 dose led to excessive immune stimulation and significant decrease in mRNA expression (Figure S3). After R848 priming, addition of LNPs further potentiated cytokine release in a manner dependent on both LNP type and dose, as measured by representative cytokines including IL-1β (Figure 2c), IL-6 (Figure 2d), and IP-10 (Figure 2e). The LNP treatment dose was capped ∼2 μg/mL of total mRNA to maintain cell viability (Figure 2f), as well as allowing the most distinctive rank ordering of cytokine responses among different LNP types (Figure 2c-e). The IL-1 cytokine responses were generally stable within day 2 to 7 after LNP treatment (Figure S4), so that cytokine analysis at day 3 post LNP treatment represented robust rank ordering results. To establish optimal R848 concentrations across different in vitro models, we performed parallel dose titrations in primary cells, including PBMCs and monocytes (Figure S5). These comprehensive titrations across all cellular models confirmed that an R848 dose of 0.1 μg/mL is optimal for differentiating mRNA-LNP-induced cytokine release from background noise. Based on these results, pre-treatment of 0.1 μg/mL R848 for 2 hrs, followed by 2 μg/mL LNP treatment for 3 days, was employed for the following iMonocyte assays.

**Figure 2:**
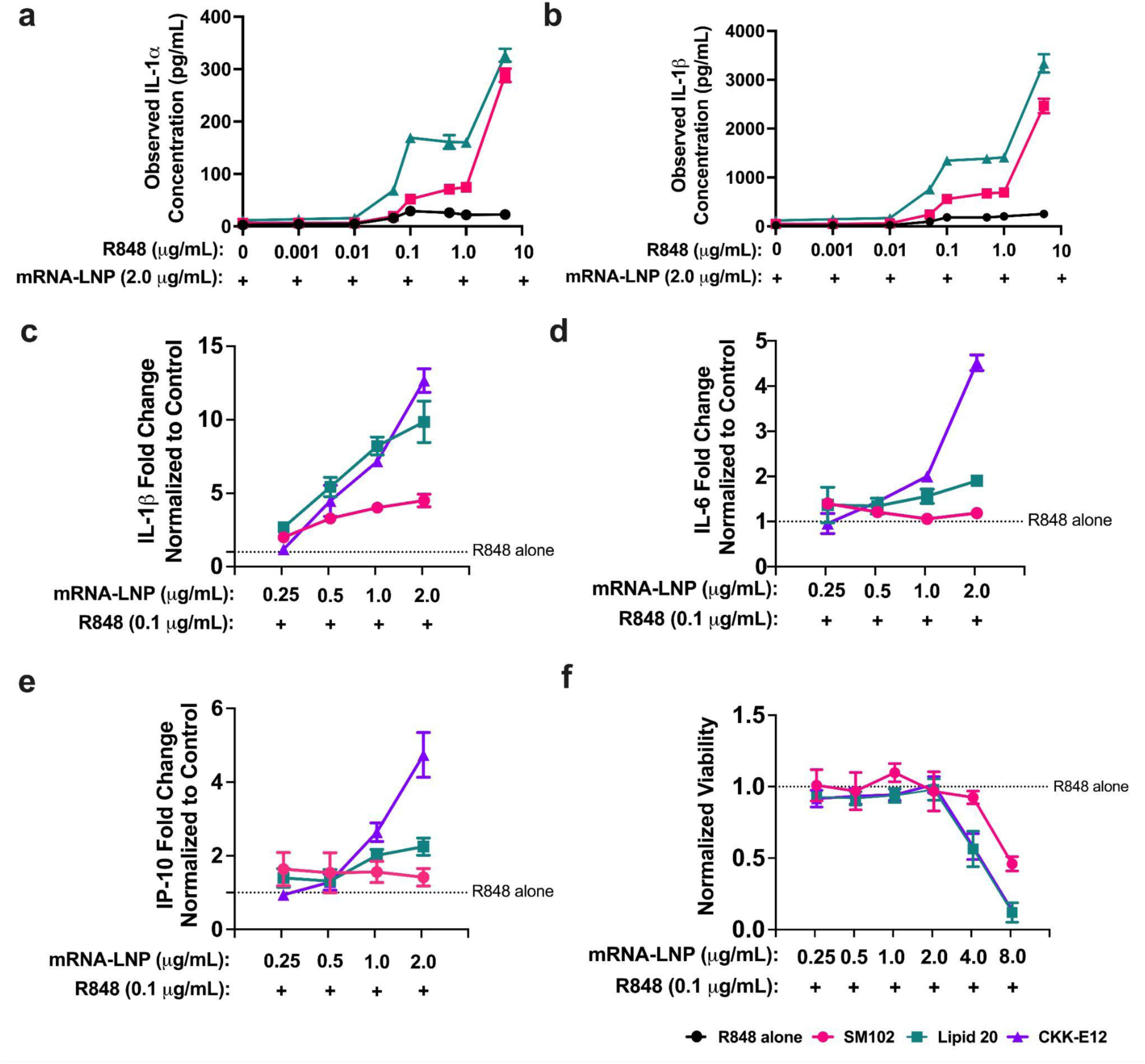
R848 priming enhances the consistency and magnitude of inflammatory responses to mRNA-LNPs in iMonocytes. **(a,b)** R848 dose titration in iMonocytes prior to mRNA-LNP treatment. Cytokine secretion was measured in conditioned media on day 3 after LNP treatment. **(c-f)** Dose-response analysis of mRNA-LNPs in iMonocytes. Cytokines (c-e) and cell viability (f) were evaluated on day 3 following mRNA-LNP treatment. Cells pre-treated with 0.1 µg/mL R848 but without mRNA-LNP treatment served as control. Results represent mean ± SEM, n = 3.

### iMonocytes improve rank ordering of LNP-induced innate immune responses compared to primary cell-based assays

Next, we compared different *in vitro* models, including primary human PBMCs and monocytes, to iMonocytes, for their performance in cytokine profiling and rank ordering of different LNPs in terms of innate immune responses (Figure 3a). As expected, donor-to-donor variability was observed in the primary immune cell models (Figure 3b). Following LNP treatment, several key cytokines, including IL-6 and CCL5 (RANTES), exhibited inverse secretion patterns between donors. For instance, in PBMCs, IL-6 fold change in response to CKK-E12 was 0.9 in Donor 1 versus 1.4 in Donor 2, complicating data interpretation (Figure 3b, S6). CCL5 showed even greater inconsistency: for SM102, Lipid 20, and CKK-E12 LNPs, CCL5 levels were generally similar to control in Donor 1 but decreased in Donor 2. Given that IL-6 and CCL5 are key mediators of innate immunity and dysregulated inflammation,^8, 29^ accurately quantifying their induction is critical. These differences likely reflect inherent physiological variation between individuals, an expected aspect of primary cell assays. As a result, ranking LNP’s innate immune responses using primary cells across different donors can present challenges. Moreover, primary immune cells lacked sufficient sensitivity to distinguish between LNPs with weak or moderate proinflammatory responses. In radar plots, although highly stimulatory LNPs like CKK-E12 triggered robust responses detectable by PBMCs or primary monocytes, responses to “weaker” LNPs overlapped substantially.

**Figure 3:**
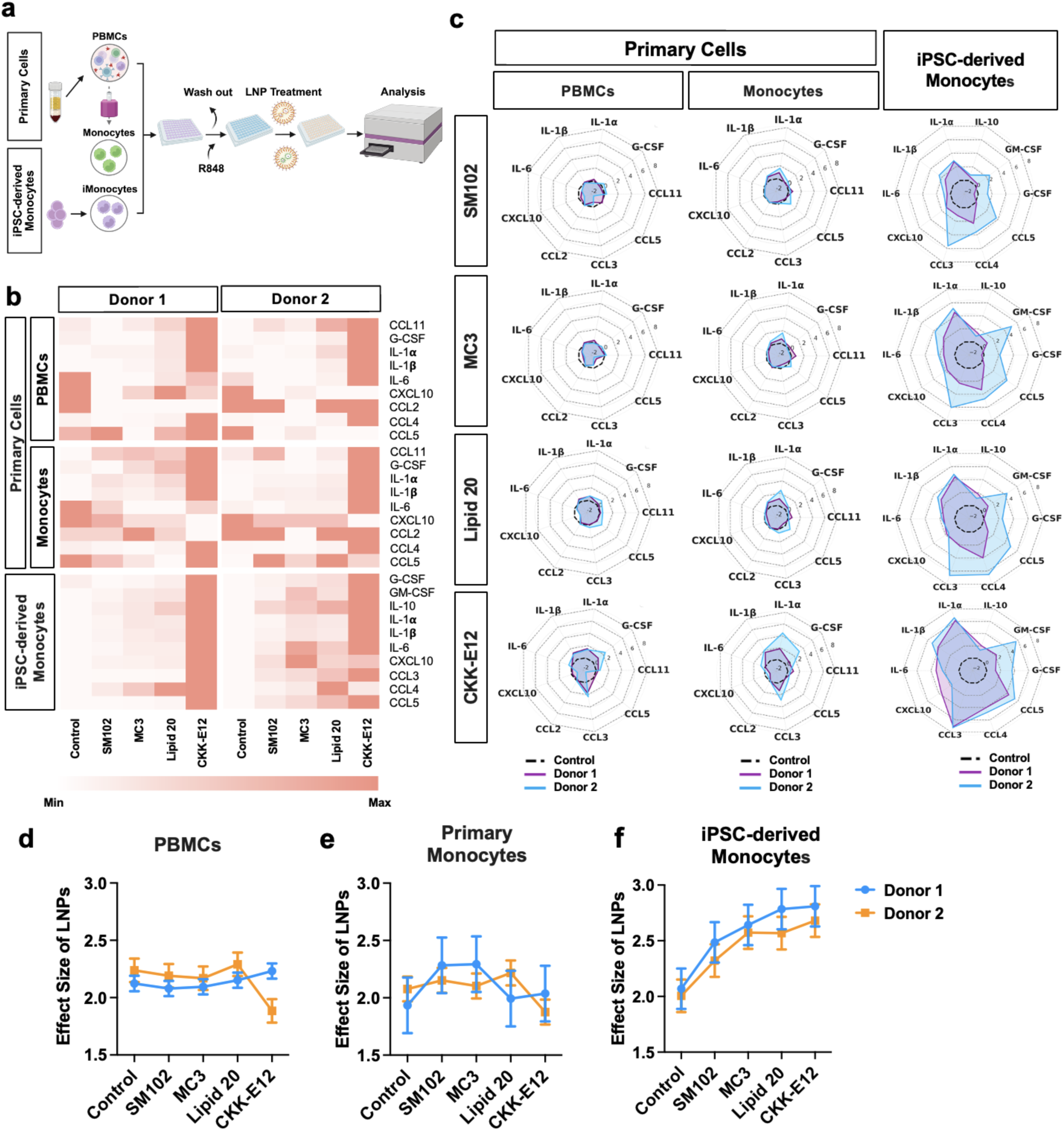
iMonocyte platform improved rank ordering of LNP-induced innate immune responses over primary PBMC and monocyte platforms. **(a)** Cell line preparation schemes for different assay platforms. **(b)** Heatmap of fold changes of LNP-induced cytokines in cell culture supernatants, relative to the R848-alone control. Primary cell lines were tested with two donors (each primary monocyte line was derived from the same donor), and iMonocyte was tested with two sources. Scale bar is adjusted according to different cytokine response ranges, with the R848 control group always set as 1. **(c)** Visualization of panel (b) data in radar plots. **(d-f)** Effect size analysis of cytokine responses for each LNP tested by different *in vitro* cell models and donors. For iMonocytes, donor 1 represents the IP11 line and donor 2 represents the SCTi003 line.

In contrast, iMonocytes demonstrated higher sensitivity and more consistent rank order of LNP-induced cytokine responses. While the SCTi003 line produced higher overall cytokine levels than the IP11 line, both preserved similar cytokine responses trending across different LNP types, as shown by the radar plots (Figure 3c). To quantitatively assess cytokine response sensitivity across the different models, we calculated the effect size of cytokine induction for PBMCs (Figure 3d) and primary monocytes (Figure 3e) from two donors and for iPSC-derived monocytes from two independent cell lines (Figure 3f). In primary cells, effect sizes clustered around 2.0, even for the highly immunostimulatory CKK-E12 LNP, indicating limited discriminatory power. By contrast, iMonocytes exhibited a clear, graded increase in effect size, effectively distinguishing low (SM102), intermediate (MC3, Lipid 20), and high (CKK-E12) immune-stimulatory LNPs. Notably, while SCTi003 and IP11 displayed differences in overall cytokine levels, which is a natural variation when comparing distinct iPSC lines,^30^ trends of their effect sizes were comparable. This consistency further underscores the reproducibility of the iMonocyte platform. This robust stratification highlights the strength of iMonocytes in sensitively and reliably assessing the immunostimulatory potential of LNPs across both donors and formulations.

Furthermore, PBMCs are primary immune cells with a limited lifespan *ex vivo*, which poses challenges for conducting longitudinal studies, such as investigating the immune response kinetics to LNPs over time or evaluating the effect of repeated dosing. By 48 hours post-isolation, PBMC populations failed to maintain normal functional parameters.^19^ In contrast, iMonocytes exhibited extended viability, with no significant cell death observed at day 7 post-harvest (Figure 1b). Additionally, within the assay time window, mRNA expression reached and remained plateau from day 2 to at least day 5, with iMonocyte viability also maintained at day 7 (Figure S7). Furthermore, mRNA expression kinetics correlated with cytokine response kinetics as measured by IL-1 (Figure S4). Taken together, these findings highlight the suitability of iMonocytes for kinetic cytokine assessments, thereby overcoming the limitations associated with short-lived, primary cell-based in vitro assays.

### iMonocyte-based cytokine profiles correlate with *in vivo* immune responses to LNPs

To further investigate the translational value of the iMonocyte assay platform, we evaluated *in vivo* cytokine responses of the panel of LNPs in a murine model (Figure 4a). Serum cytokine responses were analyzed at 6 hr after a single intramuscular (IM) injection at 0.15 mg/kg of different LNPs.

**Figure 4.**
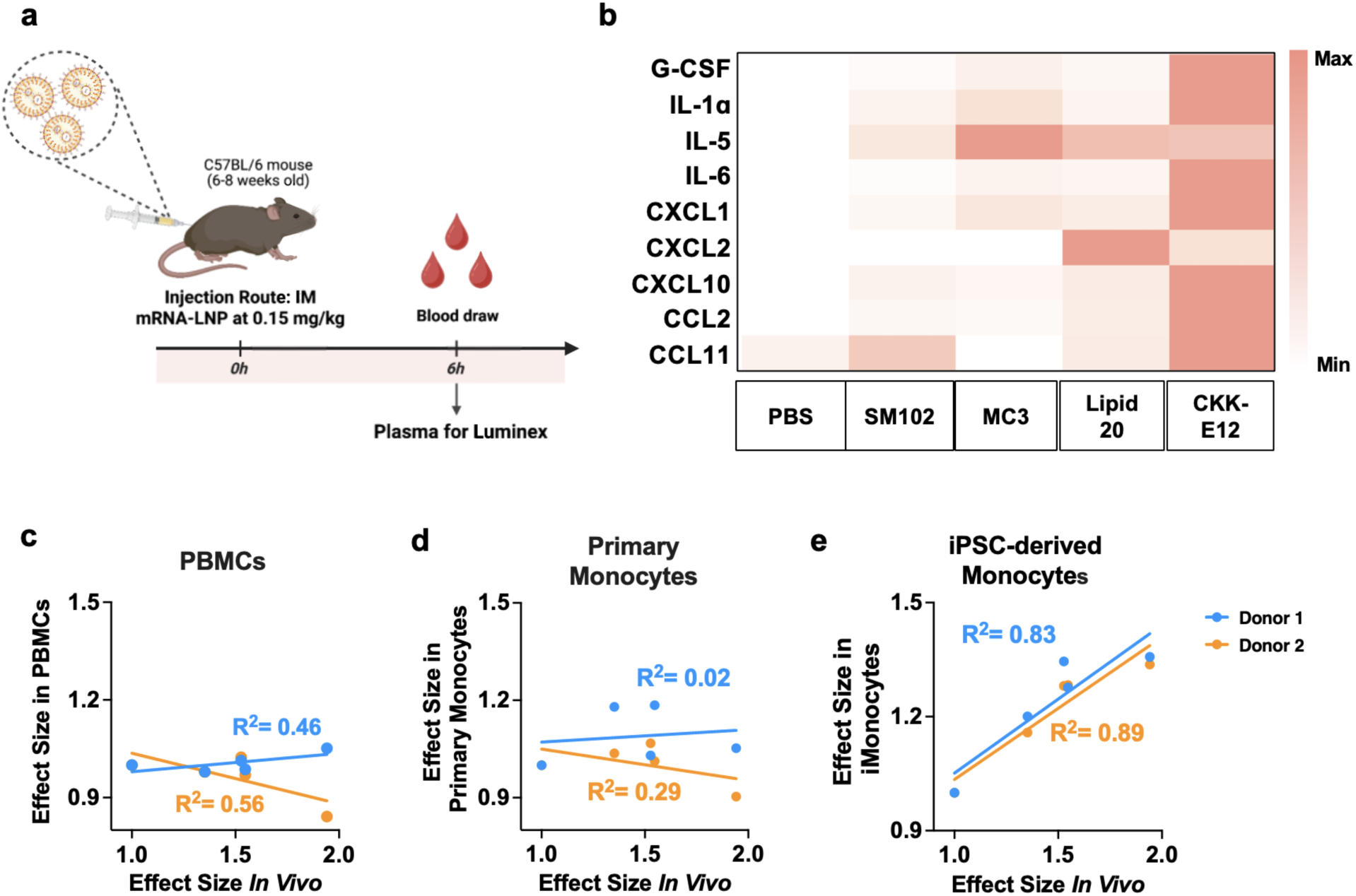
iMonocyte platform captures rank ordering of LNP-induced innate immune responses in vivo. **(a)** Experimental workflow for measuring *in vivo* cytokine response to LNPs. C57BL/6 mice (6–8 weeks old) were intramuscularly injected with 0.15 mg/kg mRNA-LNPs Blood plasma was collected at 6 hours post-injection for cytokine analysis by Luminex. **(b)** Heatmap of *in vivo* cytokine fold changes relative to the PBS control. **(c-e)** Correlations of effect sizes between *in vivo* and *in vitro* cytokine responses. iMonocyte results strongly correlated with *in vivo* results, whereas primary cells showed weak correlations.

Among the tested formulations, CKK-E12 LNP elicited the most pronounced cytokine response, while SM102 led to only modest elevations. Cytokine levels triggered by CKK-E12 exceeded those of SM102 by approximately 3- to 20-fold across several key inflammatory mediators, including IL-6, MCP-1, G-CSF, CXCL1, IL-1α, and IP-10 (Figure 4b, S8). Importantly, the moderately stimulatory LNPs, MC3 and Lipid 20, as classified by the iMonocyte assay, also induced stronger *in vivo* cytokine responses compared to SM102. This trend closely mirrored the relative immune activation profiles observed *in vitro*, supporting the translational relevance of iMonocytes in modeling *in vivo* responses.

To quantitatively assess concordance between models, we again calculated and compared the effect sizes of cytokine secretion between *in vivo* (normalized to the PBS control) and *in vitro* (normalized to the R848 control) (Figure 4c-e). iMonocytes demonstrated a strong positive correlation with *in vivo* responses (R² = 0.83 for IP11 iMonocytes and R² = 0.89 for SCTi003 iMonocytes), capturing both the magnitude and rank order of LNP immunogenicity. In contrast, primary cell-based assays exhibited poor correlation with in vivo cytokine responses, showing R² = 0.46 and 0.56 for different PBMC donors, and R² = 0.02 and 0.29 for different primary monocyte donors. The trends between different donors also showed different directions, complicating interpretation of *in vitro*-in vivo correlations.

Collectively, these findings establish the translational relevance of iMonocyte-based cytokine assays for LNPs. Within the panel of LNPs tested, the iMonocyte assay not only distinguishes their varying immunostimulatory potency *in vitro* but also correlates *in vivo* cytokine response patterns, positioning it as a promising tool for preclinical innate immune response screening.

### iMonocyte transcriptomes exhibit distinct immune profiles following different mRNA-LNP treatments

To mechanistically understand LNP-induced cytokine response, we further performed RNA-seq in the treated iMonocytes to examine their gene expression profiles in response to mRNA-LNPs with graded innate immune responses (SM102 < Lipid 20 < CKK-E12). Principal component analysis (PCA) revealed that treatment with mRNA-LNPs had a strong effect on the overall gene expression profiles, as indicated by the segregated grouping by different LNPs along the first principal component (PC1), which corresponded to the known rank ordering of immune response intensities (Figure 5a). Differential expression analysis showed that each mRNA-LNP formulation, in combination with R848, elicited robust profiles of differentially expressed genes (DEGs; |Log2FC| > 1 & FDR < 0.05) when compared with the R848-only control. The magnitude of transcriptional changes increased with the immune potency of the LNPs, in alignment with prior observations (Figure 5b, Table S1). Notably, DEGs included key cytokines such as *CCL5*, *CXCL10/IP10*, *IL1B*, and *IL1A*, which showed elevated expression levels, with high reproducibility among biological replicates within each treatment group (Figure 5c). Analysis of shared DEGs across conditions revealed substantial overlap, with 101 genes upregulated and 60 downregulated across the SM102, Lipid 20 and CKK-E12 LNP formulations (Figure 5d, Table S2). Average gene set expression scores confirmed this trend associated with the LNP immune potency ranking, as genes with increased expression revealed progressively higher scores, while the genes with decreased expression showed a lower score (Figure 5e). Gene ontology (GO) and Reactome pathway enrichment analysis of the upregulated shared DEGs highlighted functional associations with key monocyte processes, such as interferon α/β signaling and cytokine/chemokine activity, which are central to mRNA-LNP induced innate responses (Figure 5f).^10^ In addition to the gene set, Reactome pathway enrichment analyses of the individual contrast ranked interferon α/β signaling, chemokine receptors bind chemokines, and interleukin-10 signaling amongst the top significant (FDR < 0.05) pathways (Table S3). These pathways were up-regulated in all three LNP formulations and their gene set scores increased following control < SM102 < Lipid 20 < CKK-E12 LNPs, consistent with their rank ordering in innate immune responses assessed by the iMonocyte platform (Figure 5g-i). Interestingly, enrichment of the interferon α/β signaling pathway was represented mostly by interferon-stimulated genes (ISGs), such as *ISG15*, *IFIT1*, *OAS1,* whereas transcripts for Type I IFN ligands (*IFNA1*, *IFNA2, IFNB1*, *IFNB2*) were not represented (Figure S9).

**Figure 5.**
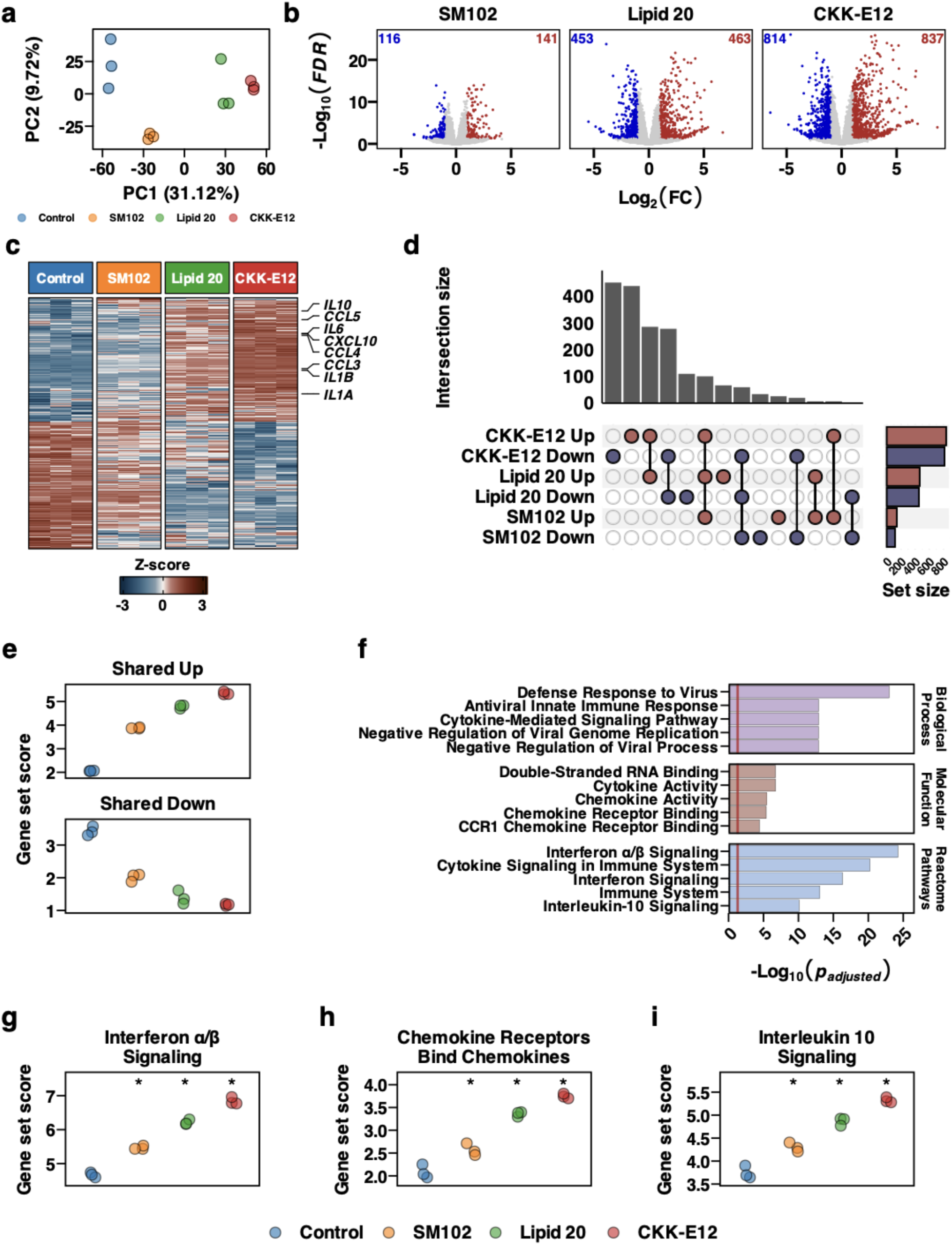
RNA-seq of R848/LNP-treated iMonocytes reveals common up-regulation and down-regulation pathways. (a) Principal component analysis (PCA) of gene expression profiles of biological replicates from different LNP treatment groups. (b) Volcano plots of differential gene expression profiles from the three mRNA-LNPs. Red and blue represent differentially expressed genes (DEGs; |Log2FC| > 1 & FDR < 0.05) with increased or decreased expression, respectively. Total numbers of DEGs for each direction are also indicated. (c) Heatmap of the Z-scores of log2CPM gene expression values from the biological replicates for all of the DEGs. (d) UpSet plot of the intersections between the DEGs with increased (red) or decreased (blue) expression. (e) Gene set scores (mean log2CPM) of the DEGs with a shared direction of change across the three LNPs, split by direction. (f) Enriched Gene Ontology (GO) categories (based on biological process and molecular function) and Reactome pathway terms for the shared genes with increased expression. **(g-i)** Gene set scores of the top up-regulated Reactome pathways from different LNP treatments, where asterisks indicate a significant (FDR < 0.05) difference when compared to R848 alone control.

Overall, the transcriptomic findings align well with the cytokine data from iMonocytes, recapitulating the immune activation rank ordering of SM102 < Lipid 20 < CKK-E12 LNPs. Collectively, these observations reinforce iMonocytes as a robust, sensitive, and mechanistically clean platform for probing LNP-induced innate immune response.

### The iMonocyte platform differentiates both lipid and mRNA cargo-induced innate immune responses

The development of mRNA-based therapeutics necessitates careful consideration of both lipid excipients and mRNA structural modifications, as these components collectively influence protein expression, innate immune activation, and therapeutic efficacy.^31, 32^ RNA engages multiple pattern recognition receptors (PRRs), including endosomal Toll-like receptors (TLR3, TLR7, and TLR8) and cytoplasmic sensors (RIG-I and MDA-5), through their characteristic sequence motifs and secondary structures.^33, 34^ Nucleoside modifications like m1Ψ offer an established approach to reducing PRR activation while maintaining translational efficiency.^35, 36^

To address this interplay, we formulated m1Ψ modified or unmodified mScarlet mRNA into SM102, MC3, Lipid20, and CKK-E12 LNPs, and assessed their effects on protein expression and cytokine secretion using the iMonocyte platform. Enhanced expression efficiency was observed for all modified mRNA-LNPs, except for Lipid 20, compared to their counterparts loaded with the unmodified mRNA (Figure 6a). This enhancement was accompanied by significantly reduced secretion of proinflammatory cytokines including IL-1β (Figure 6b) and IL-6 (Figure 6c), following SM102, MC3, or Lipid 20 LNP treatment. These results were consistent with the established role of m1Ψ in damping innate immune activation.^35, 36^ These effects were further validated using an unmodified Ovalbumin (OVA) mRNA-LNP, which elicited markedly higher cytokine levels compared to m1Ψ-modified mScarlet mRNA LNPs,^37, 38^ confirming our platform’s ability to discriminate not only LNP-triggered but also innate immune responses induced by the cargo itself (Figure S10).

**Figure 6.**
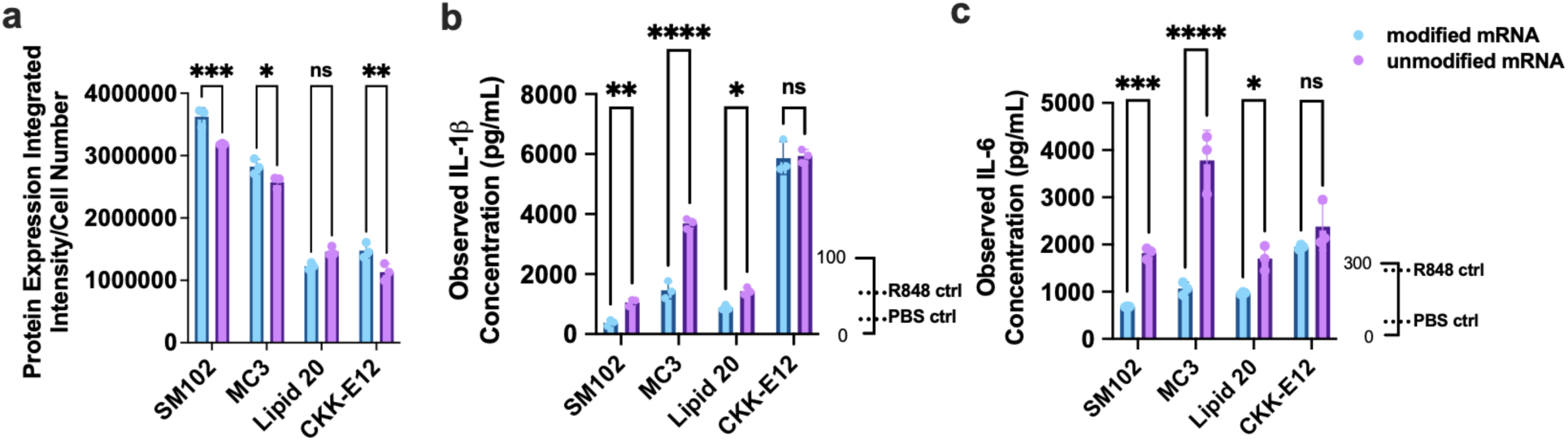
iMonocyte platform differentiates lipid- and mRNA-induced cytokine responses. **(a)** Translation efficiency of N1psU-modified versus unmodified mRNA delivered via LNPs containing different ionizable lipids (SM102, MC3, Lipid 20, CKK-E12). **(b-c)** Cytokine secretion profiles of IL-1β **(b)**, and IL-6 **(c)** following 72 hr-treatment of LNPs containing different ionizable lipids and loaded with either the modified or unmodified mRNA. Results represent mean ± SEM, n = 3. *p < 0.05, **p < 0.01, ***p < 0.001, ****p < 0.0001, as analyzed by two-way ANOVA followed by Tukey’s multiple comparisons.

The interplay between cargo and lipid types can produce distinct innate immune stimulation profiles.^8^ Analysis of cytokine responses elicited by modified versus unmodified mRNA revealed substantial variation across different LNPs. Notably, MC3 LNPs delivering unmodified mRNA induced significantly elevated levels, 2.5-fold higher IL-6, compared to the CKK-E12 LNP loaded with the unmodified mRNA (Figure 6c). Furthermore, the magnitude of cytokine level changes between modified and unmodified mRNA was greater with MC3 than with the other two LNPs tested (SM102 and Lipid 20). For CKK-E12, which is already identified as a strong innate immune stimulatory lipid, switching the cargo from modified to unmodified mRNA did not further increase the cytokine responses, probably suggesting the lipid-driven responses. These differential cytokine signatures likely reflect the engagement of different levels of innate immune signaling pathways, potentially driven by the interaction between lipid species and cargo type, which may influence how cells recognize them via TLRs and other innate immune pathways.^39^

The iMonocyte-based platform enables concurrent assessment of transfection efficiency and innate immune activation across diverse LNPs and mRNA cargos, offering a powerful tool for the rational design of next-generation mRNA therapeutics. Future applications integrating iMonocyte screening with *in vivo* immunization data could guide the development of mRNA-LNP formulations with optimized immune signatures, leading to the next generation therapeutic strategies.

### The iMonocyte platform enables evaluation of mRNA-LNP dosing regimens

Dosing regimens play a crucial role in the development of mRNA-LNP therapeutics, balancing therapeutic efficacy with safety and tolerability. While multiple approaches have been explored to understand the impact of dosing regimens on innate immune response,^40, 41^ there remains an unmet need for translational *in vitro* models to systematically evaluate these dynamics. To address this gap, we leveraged the iMonocytes which offer a prolonged testing window, to assess cytokine responses to mRNA-LNPs under different dosing schedules and total doses. Specifically, iMonocytes were treated with m1Ψ-modified mScarlet mRNA-LNPs formulated with different ionizable lipids (SM102, MC3, Lipid 20, and CKK-E12) and three dosing strategies: single dosing (1 µg/mL on day 0), split dosing (0.5 µg/mL on day 0 and 0.5 µg/mL on day 1), and repeated dosing (1 µg/mL on day 0 and 1 µg/mL on day 1), followed by cytokine analysis on day 3 (Figure 7a). Overall, split dosing at a total dose of 1 µg/mL led to elevated cytokine levels compared to single dosing across all LNP formulations tested (Figure 7b). This finding aligns with published studies that split dosing elicited stronger immune responses across various vaccine platforms when delivered to multiple tissue compartments, such as skin and muscle.^42^ These results underscore the importance of the dosing schedule in stimulating an innate immune response, which potentially shapes the overall therapeutic efficacy. We further evaluated repeated dosing, a common clinical approach exemplified by COVID-19 mRNA vaccine regimens. Results showed significantly stronger cytokine responses compared to single dosing (Figure 7b), consistent with literature demonstrating that repeat administration enhances both innate and adaptive immune responses.^3^ Interestingly, when comparing split dosing at a total dose of 1 µg/mL to repeated dosing at a total dose of 2 µg/mL, we observed that the latter dose strategy further elevated cytokine release rather than reaching a plateau. These findings suggest that iMonocytes maintain sensitivity to dosing variations, providing a robust platform for detecting cytokine secretion patterns induced by varying dosing schedules as well as total doses.

**Figure 7.**
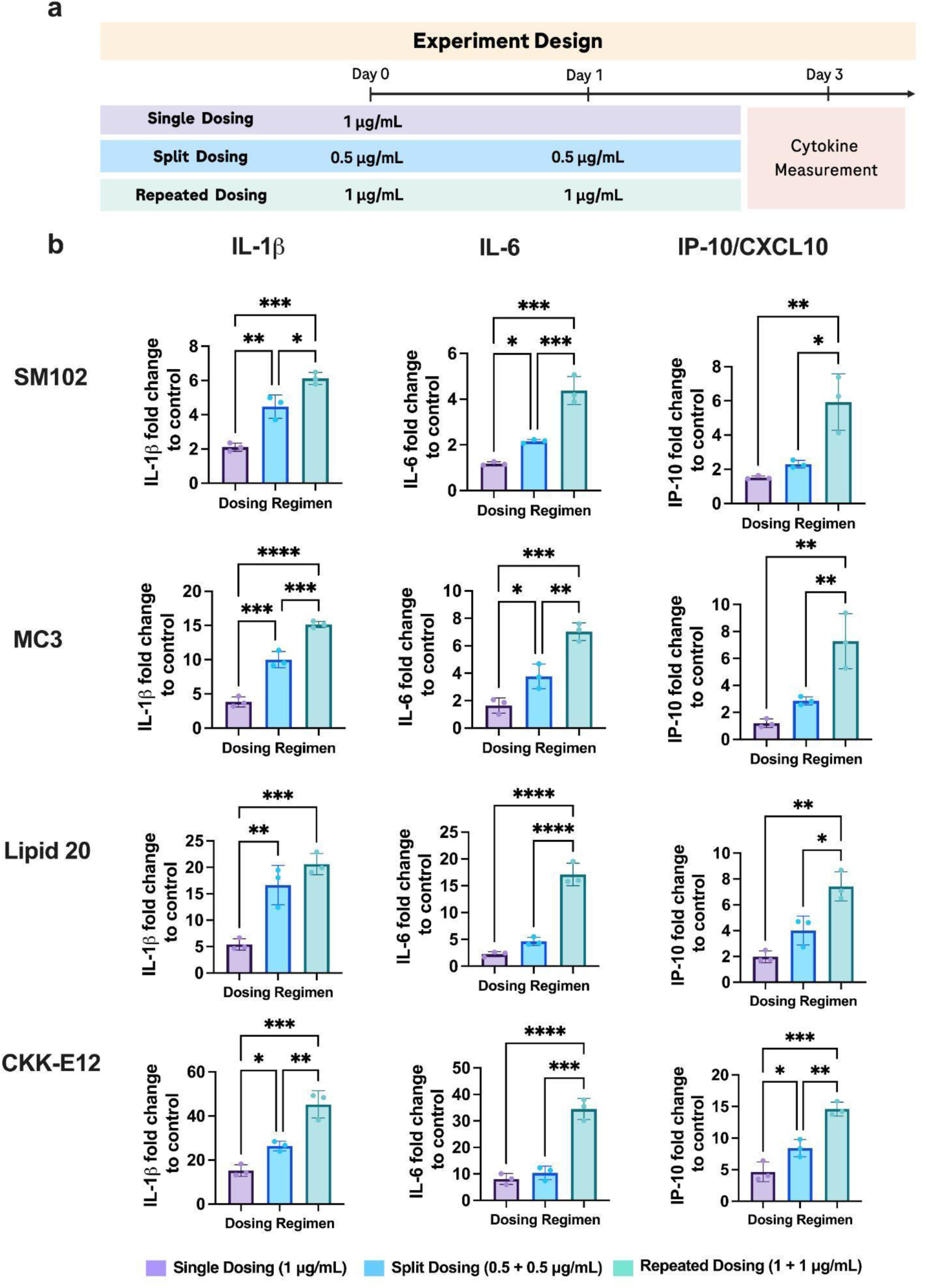
iMonocyte platform enables evaluation of innate immune responses under different LNP dosing regimens. **(a)** Experimental design for in vitro dosing regimen analysis. iMonocytes were treated with N1psU-modified mScarlet mRNA-LNPs formulated with different ionizable lipids (SM102, MC3, Lipid 20, and CKK-E12) using three dosing strategies: single dosing (1 µg/mL on day 0), split dosing (0.5 µg/mL on day 0 and 0.5 µg/mL on day 1), and repeated dosing (1 µg/mL on day 0 and 1 µg/mL on day 1). **(b)** Cytokine levels (IL-6, IP-10/CXCL10, and IL-1β) in conditioned media were measured on day 3 post-treatment using Luminex assay. Data represent mean ± SEM, n = 3 biological replicates. Statistical significance was determined by ordinary one-way ANOVA: *p < 0.05, **p < 0.01, ***p < 0.001, ****p < 0.0001.

Collectively, this study highlights the utility of the iMonocyte-based *in vitro* assay in screening mRNA-LNP immunostimulation from both ionizable lipids and payloads, as well as enabling systematic evaluation of dosing regiments. This platform offers critical insights that support the rational design of preclinical studies and could potentially inform clinical strategies to optimize mRNA–LNP-based therapies for improved safety and efficacy.

## Discussion

LNPs have revolutionized the landscape of mRNA-based therapeutics, establishing them as essential tools for delivering nucleic acids *in vivo*.^7–12^ However, their clinical potential hinges on the ability to precisely modulate innate immune activation. Traditional platforms employing primary cells, such as PBMCs, often suffer from variability in output and limited sensitivity, hindering reliable assessment of LNP immunostimulatory properties. In this study, we present iMonocytes as a scalable, reproducible, and physiologically relevant platform that bridges these limitations and offers a robust framework for evaluating LNP-induced innate immune responses.

The use of iPSCs to derive monocytes offers substantial benefits in LNP profiling. iPSCs offer an unlimited, renewable cell source, laying the foundation for personalized models to study immune responses across diverse genetic populations.^25–27^ Standardized differentiation protocols further enhance reproducibility and experimental consistency, addressing key challenges inherent to primary cell assays. While the differentiation of iPSCs into mature monocytes involves technical complexities, recent advancements in differentiation methodologies have substantially improved reliability,^27, 43^ positioning iMonocytes as an increasingly accessible system for immunological studies. Though initial setup costs may pose a barrier, their scalability, coupled with continuous progress in functional characterization, underscores the assay’s value as the next-generation tools in immunology.

In direct comparison with PBMCs and primary monocytes, iMonocytes outperformed cellular platforms by exhibiting consistent and sensitive responses to LNP formulations. Across all tested ionizable lipids and donors, iMonocytes accurately ranked immunostimulatory profiles, correlating closely with *in vivo* cytokine signatures. This predictive capability is crucial for preclinical screening, as it facilitates the early identification of overstimulatory formulations while supporting the optimization of LNP designs. Notably, mRNA modifications, such as the m1Ψ, reduced cytokine release in an ionizable lipid-dependent manner, unveiling complex interdependencies between the carrier vehicle and payload. These findings reinforce the necessity for integrated analysis of LNP composition and payload structure during screening approaches. Our results further demonstrated that iMonocytes enable systematic evaluation of dosing regimens, a critical component of preclinical development. Split and repeated dosing strategies enhanced cytokine responses compared to single dosing, aligning with *in vivo* and clinical observations from mRNA vaccines. This capability highlights the translational relevance of iMonocytes in dose optimization, reducing reliance on animal studies while accelerating the refinement of therapeutic dosing protocols.

Beyond monocyte engagement, LNPs activate the complement system - a process governed by their physicochemical properties (surface charge, PEGylation density, and lipid composition) that generates anaphylatoxins (C3a, C5a) and opsonins (C3b).^8, 44^ These complement products potentiate monocyte responses through CR3-mediated phagocytosis and proinflammatory cytokine production (IL-1β, IL-6), establishing a functional bridge between cellular innate defense mechanisms.^8, 44^ The resulting innate immune activation propagates to adaptive immunity via enhanced dendritic cell maturation and antigen presentation, culminating in robust CD8+ cytotoxic T cell responses and CD4+ T helper cell polarization.^45, 46^ Therefore, comprehensive profiling of these LNP-complement interactions via iMonocytes could further guide rational design of vaccine platforms for finely tuned immune activation profiles. Moreover, future high-throughput screening of LNP libraries may reveal LNP physicochemical structure-activity relationships that enable precise control over immunostimulatory and efficacy outcomes.

To validate the translatability of our *in vitro* findings, *in vivo* studies using murine models were conducted. While murine models offer valuable preclinical insights, and mice and humans share many conserved immune pathways, evolutionary divergence has led to key functional differences in immune recognition and response mechanisms.^47, 48^ Therefore, the results should be interpreted in parallel with human data to ensure translational relevance and a comprehensive understanding of immune responses to mRNA vaccines.

The treatment of iMonocytes with mRNA-LNPs resulted in the up-regulation of gene expression related to some of the immune pathways that we have previously observed when administering ASO-LNPs to mice via intracerebroventricular injection, specifically the interferon and related oligoadenylate synthetase (OAS) pathways.^49^ This observation suggests that part of the observed immune response may be related to cellular sensing of non-host RNA and that this response differs by LNP. Overall, the relationship between gene expression profiles and immune responses suggests that transcriptomic profiles of iMonocytes can be further developed to contribute to high-throughput screening of LNPs by providing a gene set scoring system. Comparative analysis revealed that iMonocytes capture immunostimulatory responses overlooked by primary cells, particularly for LNPs with low-to-medium immunostimulatory potency. This *in vitro* behavior closely mirrors *in vivo* observations and underscores the translational relevance of the iMonocyte assay platform. Delineating the mechanistic basis for these divergent responses will be essential for refining predictive models and advancing our understanding of LNP-triggered immune activation.

In summary, iMonocytes provide a robust and translational platform for assessing LNP innate immune response, accelerating the design of safer, more effective LNP formulations and fostering innovation across the fields of nucleic acid delivery, immunotherapy, and vaccine development.

## Methods

### iPSC culture and characterization

iPSC line iP11 and SCTi003-A were purchased from ALSTEM and Stem Cell Technology, respectively. The iP11 line was generated from the foreskin fibroblasts of a male donor, and the SCTi003 was generated from healthy female donor peripheral blood mononuclear cells (PBMCs) by each vendor. After thawing cryopreserved iPSCs, the cells were plated on Matrigel®-coated cultureware, cultured, and passaged at least three times before initiating iMonocyte differentiation. All iPSCs were maintained in mTeSR™1 media (Stemcell Technologies, #85850) and were passaged as small chunks every 3–5 days, depending on confluence, using Gentle Cell Dissociation Reagent (Stemcell Technologies, #100-0485). All iPSC lines were routinely characterized for the expression of pluripotency markers, and mycoplasma tests were performed regularly. For characterization, cells were plated in a 96-well PhenoPlate (Revvity, #6055302) and subjected to immunofluorescence staining for NANOG, OCT4, SSEA4, and TRA1-81. Total RNA was purified from cells using RNeasy Micro Kit (Qiagen, #74004) and transcribed to cDNA using the theSuperScript™ IV VILO™ Master Mix with ezDNase™ Enzyme (ThermoFisher, #11766050). Quantitative RT-PCR was performed using the QuantStudio 7 Pro system, with taqMan™ Fast Advanced Master Mix (ThermoFisher, #4444557). The following pre-designed primer sets were used (Table 1): 0.1 mL custom TaqMan Array plates (18s rRNA, OCT4, NANOG, CD68, and LYZ) (#SO 12352794) and PrimeTime™ Std qPCR Assay (KLF4, CD14, CD34 and CD48).

**Table 1.**
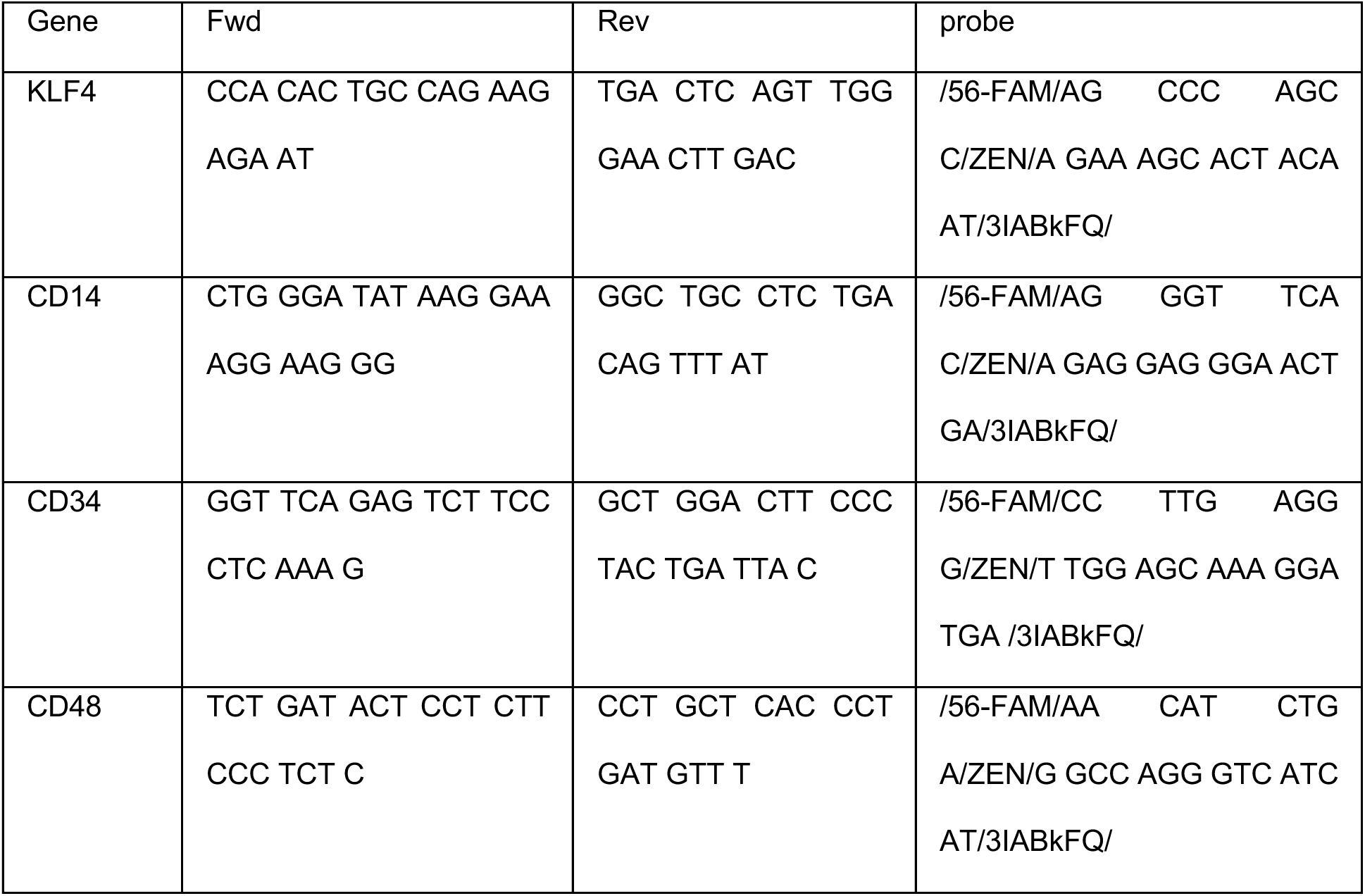
Primers and probes used for RT PCR.

### Directed differentiation of human iPSCs into iMonocyte

Hematopoietic progenitors and iMonocytes were generated sequentially from iPSCs using the STEMdiff™ Monocyte Kit (StemCell Technologies, #05320) according to the manufacturer’s instructions, with minor modifications. Briefly, iPSCs at approximately 80% confluency were dissociated using Gentle Cell Dissociation Reagent (StemCell Technologies, #100-0485) and filtered through a 37 µm reversible filter to remove single cells. The cell aggregates were then plated in Matrigel®-coated cultureware and incubated overnight at 37°C. From day 0 to day 2, the cells were maintained in Hematopoietic Medium A, followed by Hematopoietic Medium B from day 5 to day 7. A portion of the cells was collected at this stage to characterize the hematopoietic stem cells. Subsequently, Monocyte Differentiation Medium was introduced, with medium changes every 2-3 days until day 18-20, when monocytes were harvested as needed. The purity of the CD14+/CD16+ monocyte population was assessed by flow cytometry and further characterized by quantitative PCR.

### Phagocytosis assay

The uptake of pH-sensitive pHrodo dyes conjugated to zymosan A particles (P35365) by iMonocytes was assessed using high-content Incucyte imaging (Sartorius). Harvested iMonocytes were seeded into poly-D-lysine (PDL)-coated 96-well PhenoPlates (Perkin Elmer) at a density of 30,000 cells per well, one day prior to the phagocytosis assay. On the day of the assay, half of the cell culture medium was replaced with fresh medium containing 2x pHrodo particles, resulting in a final concentration of 10 µg/mL. Kinetic measurements were performed using the Incucyte system at 37°C with 5% CO2. Four images per well were captured at one-hour intervals using a 20× objective. In some wells, cytochalasin D (100 ng/mL) or LPS (200 ng/mL)was added to inhibit phagocytosis. After 48 hours of imaging, the integrated intensity (GCU × µm²/image) was calculated using the Incucyte software.

### mRNA payload synthesis

OVA mRNA and modRNA were used in the studies. OVA mRNA (SKU#L-7610) was purchased from TriLink BioTechnologies (San Diego, CA, USA). modRNA was synthesized in house, using 1-methyl-pseudouridine, capping was performed cotranscriptionally using a trinucleotide cap 1 analog (CleanCap AG (3ʹ OMe) m7(3ʹOMeG)(5’)ppp(5ʹ)(2ʹOMeA)pG, Trilink).^7^ Co-transcriptionally capped RNA reactions were performed using a Cap1 analog, CleanCap® Reagent AG (3’OMe) (TriLink, N-7413), and the T7-FlashScribe™ Transcription Kit (CellScript, Madison, WI). The linearized DNA template was mixed with ATP, CTP, GTP, m1YTP, DTT, 10× reaction buffer, RNase inhibitor, and T7 RNA polymerase. The reaction included N1-methyl pseudouridine 50-triphosphate (m1YTP) (TriLink) for nucleoside-modified mRNA, in which 100% of uridine was substituted with m1Y. For each sample, 1 µg of linearized DNA template was mixed with 2 µL of 10X T7 Transcription Buffer, 1.8 µL of each NTP (ATP, CTP, GTP, m1YTP), 1.8 µL of the CleanCap AG, 2 µL of DTT, 0.5 µL of RNase inhibitor, 2 µL of T7 FlashScribe Enzyme, and RNase-free water to reach 20 µL total. The reaction was incubated for 2 h at 37°C at 300 rpm (Thermo Scientific™ Sorvall™ Legend™ Micro 17R Microcentrifuge). DNase was then added and incubated for 15 min at 37°C. The RNA reaction was stopped by ammonium acetate precipitation. Briefly, the sample was incubated for 15 min on ice and then centrifuged at 13,300 rpm for 15 min at 4°C. The supernatants were removed, and the pellets were gently rinsed with 70% ambient ethanol. The centrifugation step was repeated under the same conditions. The ethanol supernatant was pipetted out, and the pellet was briefly air-dried prior to suspension in RNase-free water. A final column purification using the MEGAclear™ Kit (Invitrogen) was performed according to the manufacturer’s instructions. RNA concentration was measured using a NanoDrop spectrophotometer (Thermo Fisher Scientific) at 260 nm. RNA samples were stored at -80°C.

### mRNA-LNP preparation

Lipids, including 1,2-distearoyl-sn-glycero-3-phosphoethanolamine (DOPE), 1,2-dimyristoyl-sn-glycero-3-phosphoethanolamine-N-[methoxy(polyethylene glycol)-2000] (14:0 PEG2000 PE) were purchased from Avanti Polar Lipids (Alabaster, AL, USA). The ionizable lipids dilinoleylmethyl-4-dimethylaminobutyrate (DLin-MC3-DMA) (MC3); heptadecan-9-yl 8-((2-hydroxyethyl) (6-oxo-6- (undecyloxy)hexyl)amino)octanoate ester (SM102); and 3,6-bis(4-(bis(2-hydroxydodecyl)amino)butyl) piperazine-2,5-dione (CKK-E12) were from MedChemExpress (MCE, NJ, USA), and cholesterol was sourced from Sigma-Aldrich (St. Louis, MO, USA). Lipid 20, described in US patent US10,221,127, was synthesized in-house.

mRNA was diluted in a citrate buffer (50 mM, pH 3.0). The organic phase consisted of ionizable, DOPE, cholesterol, and C14-PEG2000 in ethanol, with a molar ratio of 50:10:38.5:1.5 and a total lipid concentration of 12.5 mM. mRNA-LNPs were formulated by the microfluidic method, as previously described.^50^ Specifically, NanoAssemblr Ignite (Precision Nanosystems, BC, Canada) was used to prepare bench-scale LNPs, with a total flow rate of 12 mL/min and an aqueous to organic ratios of 3:1. The resulting mRNA-LNPs were dialyzed against phosphate-buffered saline (PBS, pH 7.4) using the Pierce microdialysis plate (Thermofisher) with a 10 kDa molecular weight cutoff (MWCO). Dialysis was performed for 2 hrs at room temperature, followed by buffer change and then overnight dialysis at 4°C.

### Physicochemical Characterization of mRNA-LNPs

Particle size distributions of mRNA-loaded LNPs were analyzed by dynamic light scattering (DLS). Samples were diluted 10-fold in phosphate-buffered saline (PBS, pH 7.4) and analyzed in a 384-well microplate (Corning 3821BC, NY, USA) using a DynaPro Plate Reader III (Wyatt Technology, CA, USA). The mean particle diameters and particle size distributions, represented as percent polydispersity (%PD), were determined. The concentrations of soluble and total mRNA (after extraction using 0.2% Triton X-100 in 1× TE buffer) were quantified using the Quant-iT™ RiboGreen reagent. The percent encapsulation efficiency (%EE) of mRNA was calculated using the following equation:

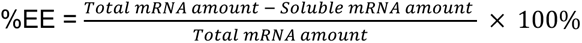

### Cryogenic transmission electron microscopy (Cryo-TEM)

Cryo-EM samples were prepared by applying 3 μL of undiluted mRNA-LNP onto glow-discharged carbon grids (Quantifoil Cu 300 mesh R1.2/1.3) treated for 7-9 s at 20 mA, followed by vitrification using a Vitrobot Mark IV system maintained at 4°C and 95% humidity with the following blotting parameters: -12 to -25 blot force, 45 s wait time, and 0.5 s drain time. Images were taken on a Talos Arctica 200 keV TEM equipped with a Falcon III detector, collecting data at 0.78-0.96 Å/pixel with defocus values ranging from -1.5 to -3 μm and a total electron dose of 60-64 e-/Å².

### PBMC isolation and primary monocyte purification

Whole blood was obtained from three to six healthy human donors who voluntarily consented to participate in Genentech’s *Samples for Science* blood donation program. The collected blood was diluted 1:1 with phosphate-buffered saline (PBS) and processed for peripheral blood mononuclear cell (PBMC) isolation using density gradient centrifugation (1200 × g for 15 minutes) in SepMate™ tubes (StemCell Technologies, Vancouver, Canada). Mononuclear cells were carefully collected from the plasma/Ficoll interface using a serological pipette.

The isolated PBMCs were washed with 1× PBS and passed through a 70 µm cell strainer to remove debris. Primary monocytes were subsequently purified from the PBMCs using the Monocyte Isolation Kit (StemCell Technologies) following the manufacturer’s protocol. Live cell counts were determined using a Vi-CELL BLU Cell Viability Analyzer (Beckman Coulter, Brea, CA, USA). PBMCs or monocytes were then seeded in a 96-well round-bottom plate at a density of 2-4 × 10⁵ cells per well and cultured in RPMI-1640 medium supplemented with 10% fetal bovine serum (FBS), 1× GlutaMAX™, 55 µM β-mercaptoethanol, 10 mM HEPES, 1× non-essential amino acids, and 1× sodium pyruvate.

### Flow cytometry

Media were removed, and cells were washed twice with PBS before staining. Viability staining was performed using Ghost Dye UV 450 (Cytek, USA) and Human TruStain FcX block (BioLegend, CA, USA) for 30 minutes at 4 °C. After staining, cells were washed and resuspended in 30 µL of antibody cocktail per well. Cells were stained for surface markers for 30 minutes at 4 °C using the following antibodies: BB700 anti-CD14 (M5E2) and PE anti-CD16 (3G8). All antibodies were obtained from either BD Biosciences (CA, USA) or BioLegend (CA, USA). Cells were acquired on a CytoFLEX LX 6-laser flow cytometer equipped with CytExpert Software (Beckman Coulter, CA, USA). Data analysis was performed using FlowJo software (v.10.8.2). Dead cells were excluded based on viability dye staining, and single cells were gated using forward scatter area and height (FSC-A/FSC-H) parameters.

### TLR agonist and mRNA-LNP treatments

The TLR7/8 agonist R848 was used to prime iMonocytes and PBMCs. A dose titration of R848, ranging from 0.01 µg/mL to 5 µg/mL, was conducted separately on iMonocytes, PBMCs, and primary monocytes to determine the optimal concentration. Cells were treated with R848 and incubated for 2 hrs at 37 °C. Cells were washed out before mRNA-LNP treatment.

mRNA-loaded LNPs were diluted 1:10 or 1:20 in Monocyte Differentiation Medium and added to cells at the final mRNA concentration of 1 or 2 µg/mL in a total volume of 200 µL per well. After overnight incubation and a subsequent 3-day culture, the plates were centrifuged, conditioned media were collected, and cells were resuspended in FACS staining buffer (1× PBS with 2% BSA) for flow cytometry analysis. Alternatively, cells were seeded into 96-well black/clear flat-bottom plates to measure mRNA expression kinetics by imaging mScarlet fluorescence intensity (Ex/Em 569 nm/594 nm) using the Incucyte (Sartorius).

### *In vivo* studies

To measure the cytokine level *in vivo*, 6–8-week-old C57BL/6 mice (Jackson Laboratory) were intramuscular injections of m1Ψ-modified mRNA–LNPs at a total mRNA dose of 0.15 mg/kg. At 6 h post-injection, whole blood was collected into tubes containing 3.8% sodium citrate as an anticoagulant. Samples were centrifuged at 1,500 × g for 10 min at 4 °C to obtain plasma, which was immediately stored at −80 °C until subsequent Luminex cytokine analysis. All animal studies were reviewed and approved by Genentech’s Institutional Animal Care and Use Committee.

### Luminex assay

To measure cytokines, conditioned media from cell culture were collected and frozen at -80 °C until the assay. Samples were thawed and analyzed using Milliplex MAP reagents (MilliporeSigma, St. Louis, MO) according to the manufacturer’s recommended protocol. The reconstituted standards were diluted by 2.73 folds to increase the number of data points from six to nine while maintaining the original dynamic range. Fluorescence intensity was measured by xPONENT software v 4.2 on FlexMap 3D instruments (Luminex Corp, Austin, TX). Standards were prepared with two replicates and measured for median fluorescence intensity using the Bio-Plex Manager v 6.2 (Bio-Rad Laboratories, Hercules, CA). Four- or five-point standard curves were obtained for each cytokine analyte and each plate. Sample measurements were performed with three replicates. Any values below the quantification limit were recorded as zero.

### RNA-seq

Total RNA was purified from cells using RNeasy Micro Kit (Qiagen, #74004) and quantified with the Quant-iT RiboGreen RNA Kit (Thermo Fisher Scientific) on a Victor X2 Multilabel Microplate Reader (PerkinElmer). RNA quality was determined using the High Sensitivity RNA ScreenTape Assay on a TapeStation 4200 (Agilent Technologies). Sequencing libraries were prepared using the SMART-Seq Total RNA PicoInput (ZapR Mammalian) kit (Takara Bio) with 1-10 nanograms of total RNA as input. Libraries quantification was performed using the Quant-iT PicoGreen dsDNA Assay Kit (Thermo Fisher Scientific) on the Victor X2 Multilabel Microplate Reader (PerkinElmer). Average library size was analyzed using D5000 ScreenTape Assay on a TapeStation 4200 (Agilent Technologies). Libraries were pooled and sequenced on NovaSeq X Plus (Illumina) to generate 30 million single-end, 50 base pair reads for each sample. RNA-seq data was analyzed through fastp,^51, 52^ GSNAP,^51–53^ HTSeqGenie,^54^ and voom-limma.^55^ Enrichr was utilized for the GO and Reactome analysis of the shared DEGs.^56–58^ CAMERA^59^ was utilized for enrichment analysis of Reactome pathways for the individual contrasts^60^.

### Statistical analysis

All quantitative data are expressed as mean ± standard error of the mean (SEM), with exact sample sizes (n) detailed in respective figure legends. Parametric analyses were performed using unpaired Student’s *t*-tests for pairwise comparisons or one-/two-way ANOVA for multi-group evaluations, followed by Tukey’s post hoc tests where appropriate. A significance threshold of p < 0.05 was applied for all analyses, which were analyzed using GraphPad Prism 10 (GraphPad Software). Cytokine induction was analyzed using fixed-effects linear regression models implemented in R (version 4.3.2). To normalize the distribution of cytokine induction levels and stabilize variance, all measurements were log-transformed using the formula log₁₀(x+1). Across all systems, treatment effects were estimated directly, with models fitted separately for the relevant experimental unit (donor or iPSC line). Replicates exhibited minimal variance and were therefore averaged prior to modeling. To ensure valid cross-system comparisons, the number of cytokines included in the models was matched between datasets: iPSC-derived monocyte analyses were restricted to cytokines also measured in the corresponding primary cell experiments, and in vivo analyses were limited to cytokines measured in vivo. For in vivo datasets, cytokine responses were modeled to obtain direct estimates of cytokine induction for each LNP formulation. For data derived from primary human PBMCs and monocytes, cytokine responses were analyzed independently for each donor, with models fitted separately for each cell type–donor combination. For iPSC-derived monocyte datasets, cytokine responses were modeled while adjusting for batch effects to control for systematic differences across experimental days. Models were fitted separately for each iPSC-derived monocyte line, with replicates averaged prior to modeling. Estimated coefficients for the treatment effects were extracted from each model. These coefficients were then aggregated across all common cytokines to estimate a composite effect size.

## Acknowledgments

We thank the Genentech core facility teams, including Flow Core, next-generation sequencing (NGS) and gCell, for their invaluable support with the Luminex cytokine assay, bulk RNA sequencing, and cell maintenance and distribution, respectively; Mercedesz Balaz for supporting this work with helpful discussion, review and feedback provided; Archana Chavan for review and providing feedback. We also appreciate the editorial comments of Karthik Nagapudi and Rajita Pappu.

## Author contributions

B.X., S.K., Y.F., C.Y., and M.E. were responsible for the conceptualization and study design, as well as contributing to the original draft writing. S.T. contributed advice on specific aspects of the study design. S.K., B.X., B.L. and Y.G. conducted the investigation and formal analysis. Data curation was handled by S.K., B.X., B.L., E.T., D.M., M.H, P.J., and D.M. K.H. and E.F. provided the materials. Technical support was given by A.A., D.R., R.L., and G.R. Finally, all authors participated in the writing – review & editing of the manuscript.

## Materials & Correspondence

Correspondence and requests for materials should be addressed to Yuchen Fan, Chun-Wan Yen, or Maheswara Reddy Emani.

## Data availability

All data supporting this study’s findings are included in the manuscript and are available from the authors upon reasonable request. Source data is provided in this paper. RNA-seq data has been deposited in GEO (accession number will be provided upon acceptance).

**AI use:** During the preparation of this work the author(s) used Google Gemini and ChatGPT in order to polish the language. After using this tool/service, the author(s) reviewed and edited the content as needed and take(s) full responsibility for the content of the publication.

