## Supplementary Figures for "Predictive *in vitro* profiling of LNP-induced innate immune response using an iPSC-derived monocyte model"

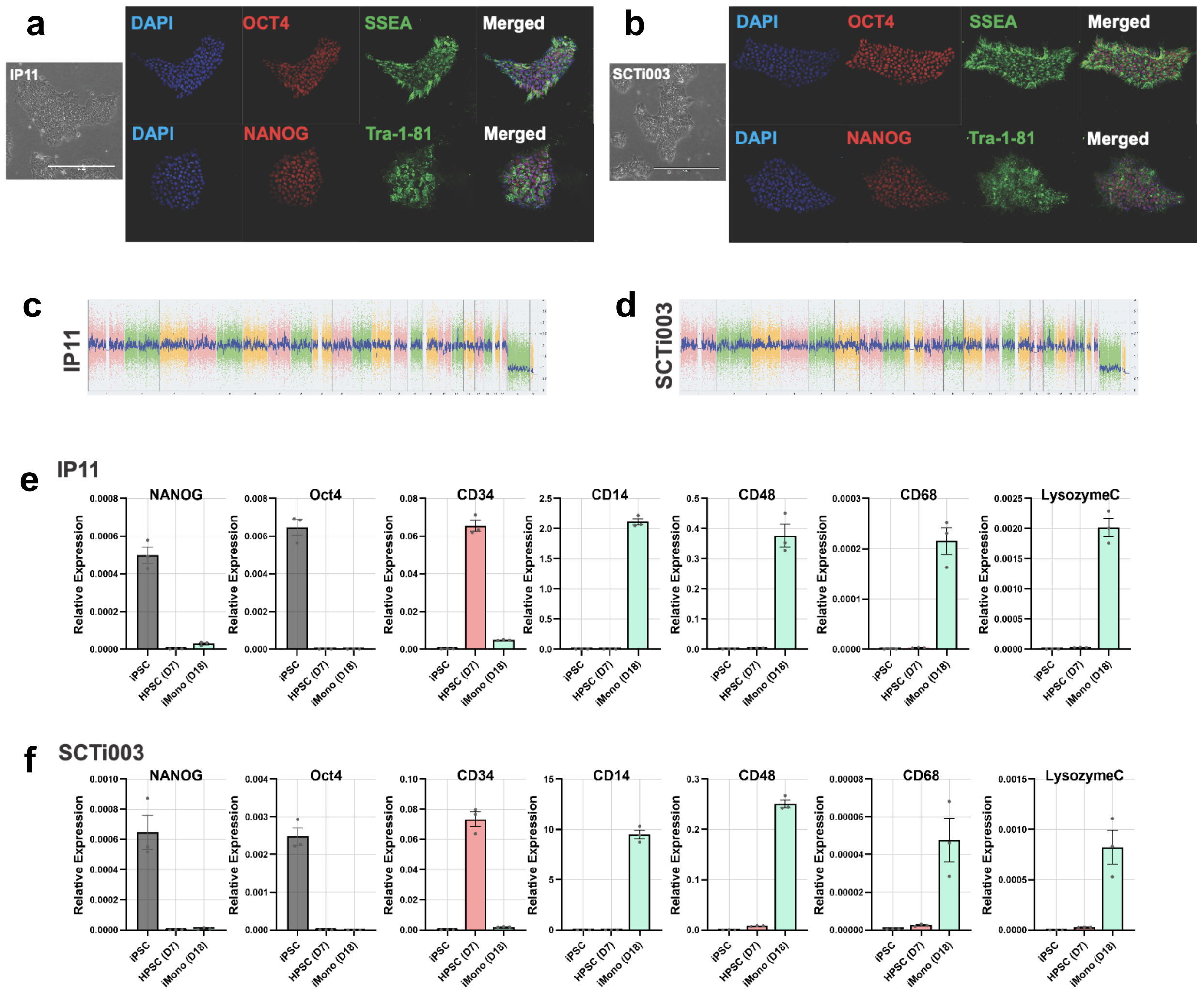


**Supplementary Figure 1. Characterization of iPSCs and iPSC-derived monocytes.**

**(a-b)** Immunocytochemical analysis of **(a)** IP11 and **(b)** SCTi003 iPSCs, demonstrating expression of pluripotency markers Oct4, SSEA1, TRA-1-81, and Nanog. **(c-d)** Karyostat analysis of **(c)** IP11 and **(d)** SCTi003 iPSCs, confirming normal karyotypes with no detectable chromosomal aberrations. Whole genome views display copy number variations across all somatic and sex chromosomes. The smoothed signal plot (right y-axis) represents the log2 ratios of microarray probe intensities, with individual chromosome probe signals indicated by pink, green, and yellow, and normalized probe signals by blue. **(e-f)** Quantitative analysis of marker gene expression in iPSCs, HPSCs at day 7, and iMonocytes at day 18. Relative expression values are normalized to GAPDH. iPSC markers (gray): NANOG, OCT4; HPSC marker (pink): CD34; monocyte markers (green): CD14, CD48, CD68, LYZ. Results are from n = 3 for each cell type.

**a**

| Sample ID | Diameter (nm) | PDI | Encapsulation Efficiency% |
|---|---|---|---|
| SM102 | 123 ± 11 | 0.18 | 97.72% |
| MC3 | 125 ± 8 | 0.15 | 96.14% |
| Lipid 20 | 118 ± 10 | 0.19 | 98.53% |
| CKK-E12 | 138 ± 15 | 0.20 | 84.44% |

**b**

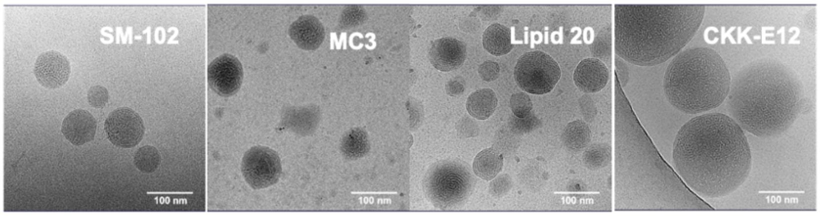


**Supplementary Figure 2. LNP formulation characterizations.**

**(a)** Particle size distribution and mRNA encapsulation efficiency of the tested LNPs. **(b)** cryo-TEM images of different LNP formulations. Scale bar in cryo-TEM images, 100 nm.

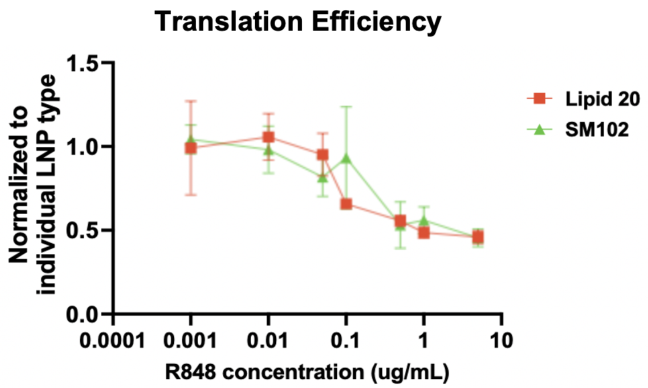


**Supplementary Figure 3. mRNA translation in iMonocytes under different pre-treatment doses of R848.**

Fold changes of translation efficiency of SM102 and Lipid 20 mRNA-LNPs in iMonocytes pre-treated with varying doses of R848 (0-8 μg/mL), followed by treatment with 2 μg/mL mScarlet mRNA-LNPs. Translation efficiency was assessed on day 3 by Incucyte imaging of mScarlet fluorescence and normalized by individual LNP type. Results represent mean ± SEM, n = 3.

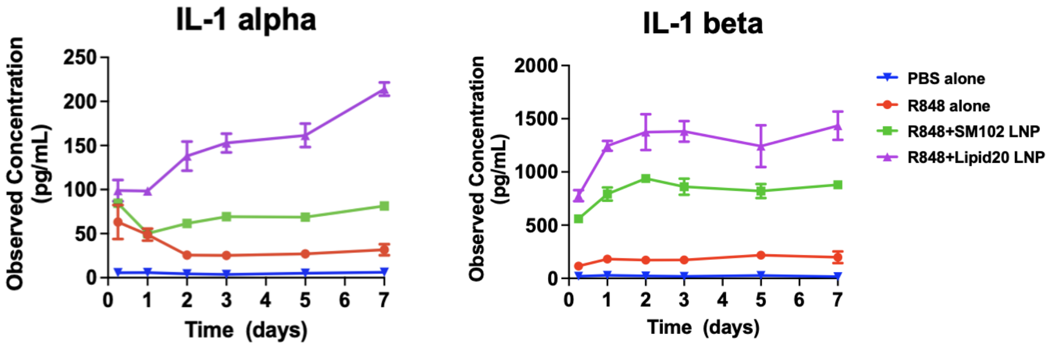


**Supplementary Figure 4. Kinetics of cytokine secretion from iMonocytes.**

IL-1α and IL-1β levels in iMonocytes from day 0 to day 7 after treatment with SM102 or Lipid 20 mRNA-LNPs. LNP-treated groups were pre-treated with 0.1 μg/mL R848. Data represent mean ± SEM, n = 3.

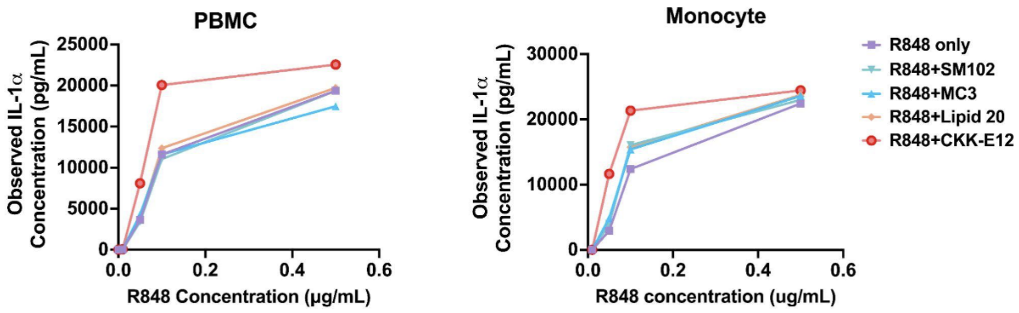


**Supplementary Figure 5. R848 dose titration in primary cells.**

PBMCs and PBMC-derived monocytes were pre-treated with different doses of R848, followed by 2 µg/mL LNP treatment for 3 days, and measured for secretion of IL-1 cytokines. Results represent mean ± SEM, n = 3.

| | | Donor 1 | | | | | Donor 2 | | | | |
|---|---|---|---|---|---|---|---|---|---|---|---|
| | | Control | SM102 | MC3 | Lipid 20 | CKK-E12 | Control | SM102 | MC3 | Lipid 20 | CKK-E12 |
| Primary Cells – PBMCs | CCL11 | 1.0 | 0.9 | 1.0 | 1.1 | 1.3 | 1.0 | 1.2 | 1.1 | 1.4 | 1.5 |
| | G-CSF | 1.0 | 0.8 | 0.8 | 1.2 | 2.0 | 1.0 | 1.3 | 1.0 | 2.0 | 5.5 |
| | IL-1α | 1.0 | 1.2 | 1.2 | 1.6 | 2.8 | 1.0 | 0.8 | 0.9 | 1.6 | 2.6 |
| | IL-1β | 1.0 | 1.0 | 1.1 | 1.4 | 2.3 | 1.0 | 1.0 | 1.0 | 1.2 | 2.0 |
| | IL-6 | 1.0 | 0.7 | 0.7 | 0.8 | 0.9 | 1.0 | 0.7 | 0.7 | 0.8 | 1.4 |
| | CXCL10 | 1.0 | 0.8 | 0.9 | 1.0 | 0.9 | 1.0 | 0.5 | 0.6 | 0.5 | 0.5 |
| | CCL2 | 1.0 | 0.9 | 0.9 | 0.9 | 0.9 | 1.0 | 1.0 | 0.9 | 1.0 | 1.0 |
| | CCL4 | 1.0 | 0.6 | 0.7 | 0.9 | 3.3 | 1.0 | 0.8 | 0.7 | 1.3 | 5.0 |
| | CCL5 | 1.0 | 1.1 | 0.7 | 1.0 | 1.1 | 1.0 | 0.3 | 0.5 | 0.3 | 0.3 |
| Primary Cells – Monocytes | CCL11 | 1.0 | 1.5 | 1.6 | 1.5 | 1.9 | 1.0 | 1.2 | 0.9 | 1.0 | 1.3 |
| | G-CSF | 1.0 | 0.9 | 1.2 | 1.4 | 1.6 | 1.0 | 1.3 | 0.9 | 1.5 | 8.7 |
| | IL-1α | 1.0 | 1.9 | 1.6 | 2.1 | 3.5 | 1.0 | 3.1 | 3.1 | 5.6 | 22.2 |
| | IL-1β | 1.0 | 1.5 | 1.3 | 1.7 | 3.1 | 1.0 | 1.6 | 1.5 | 1.8 | 4.0 |
| | IL-6 | 1.0 | 0.9 | 0.8 | 0.8 | 0.9 | 1.0 | 0.9 | 1.0 | 1.0 | 1.6 |
| | CXCL10 | 1.0 | 0.9 | 0.7 | 0.7 | 0.5 | 1.0 | 0.9 | 0.9 | 0.9 | 0.7 |
| | CCL2 | 1.0 | 1.0 | 1.1 | 1.0 | 0.9 | 1.0 | 1.0 | 0.9 | 1.0 | 1.0 |
| | CCL4 | 1.0 | 0.8 | 0.7 | 0.9 | 2.4 | 1.0 | 0.9 | 0.8 | 1.5 | 7.3 |
| | CCL5 | 1.0 | 0.9 | 0.8 | 0.7 | 1.0 | 1.0 | 2.0 | 1.5 | 2.0 | 1.5 |
| iPSC-derived Monocytes | G-CSF | 1.0 | 0.7 | 0.8 | 1.0 | 2.4 | 1.0 | 2.5 | 4.4 | 5.6 | 12.4 |
| | GM-CSF | 1.0 | 1.3 | 2.0 | 2.2 | 4.5 | 1.0 | 5.6 | 33.7 | 19.8 | 50.2 |
| | IL-10 | 1.0 | 1.5 | 2.0 | 2.7 | 4.1 | 1.0 | 1.7 | 1.9 | 2.2 | 2.5 |
| | IL-1α | 1.0 | 6.8 | 23.2 | 21.5 | 71.1 | 1.0 | 7.5 | 35.2 | 27.8 | 96.6 |
| | IL-1β | 1.0 | 2.6 | 5.0 | 6.6 | 18.1 | 1.0 | 5.5 | 13.4 | 10.0 | 37.0 |
| | IL-6 | 1.0 | 1.4 | 2.8 | 2.8 | 9.6 | 1.0 | 1.9 | 4.6 | 3.3 | 5.1 |
| | CXCL10 | 1.0 | 2.3 | 2.7 | 3.5 | 12.2 | 1.0 | 2.4 | 4.6 | 2.8 | 3.3 |
| | CCL3 | 1.0 | 2.2 | 2.5 | 4.5 | 137.0 | 1.0 | 49.4 | 65.7 | 101.7 | 154.3 |
| | CCL4 | 1.0 | 5.0 | 10.0 | 15.8 | 16.4 | 1.0 | 11.9 | 26.3 | 93.2 | 27.3 |
| | CCL5 | 1.0 | 1.1 | 1.4 | 2.2 | 20.6 | 1.0 | 13.0 | 19.6 | 30.1 | 42.9 |

**Supplementary Figure 6. Heatmap of cytokine fold change values corresponding to the heatmap shown in Figure 3b.**

Primary cells (PBMCs and monocytes) and iMonocytes were tested with two donors or sources. For iMonocytes, donor 1 represents the IP11 line and donor 2 represents the SCTi003 line.

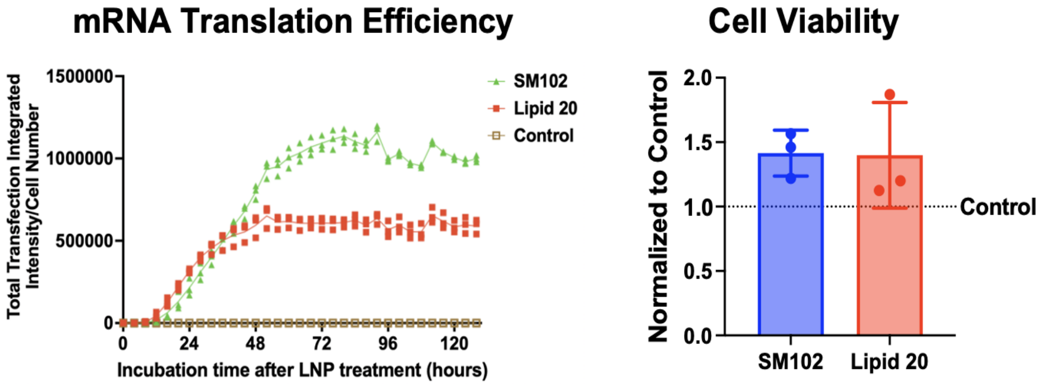


**Supplementary Figure 7. mRNA expression kinetics and viability of iMonocytes.**

iMonocytes were primed with 0.1 µg/mL R848 followed by 2 µg/mL mRNA-LNP treatment. Kinetics of mRNA expression was monitored by mScarlet fluorescence intensity within 5 days post LNP treatment. Cell viability was measured at day 7 post LNP treatment. Data represent mean ± SEM, n = 3.

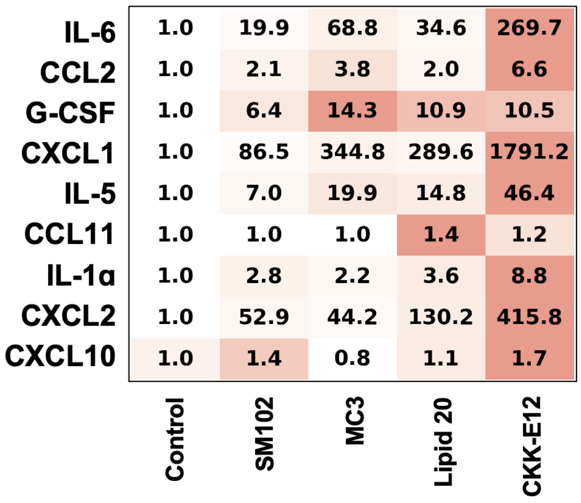


**Supplementary Figure 8. Heatmap of cytokine fold change values corresponding to the heatmap shown in Figure 4b.**

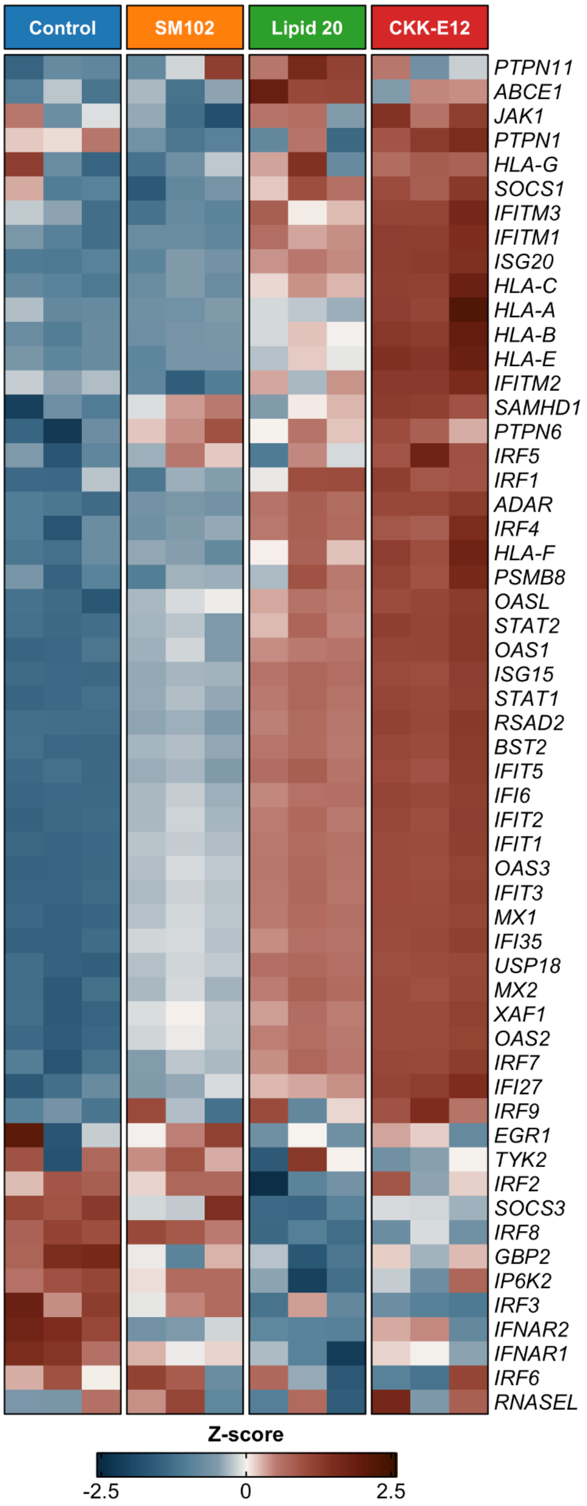


**Supplementary Figure 9. Heatmap of the Z-scores of $log_2$CPM gene expression values from the biological replicates for the detected genes from the Reactome interferon α/β signaling pathway.**

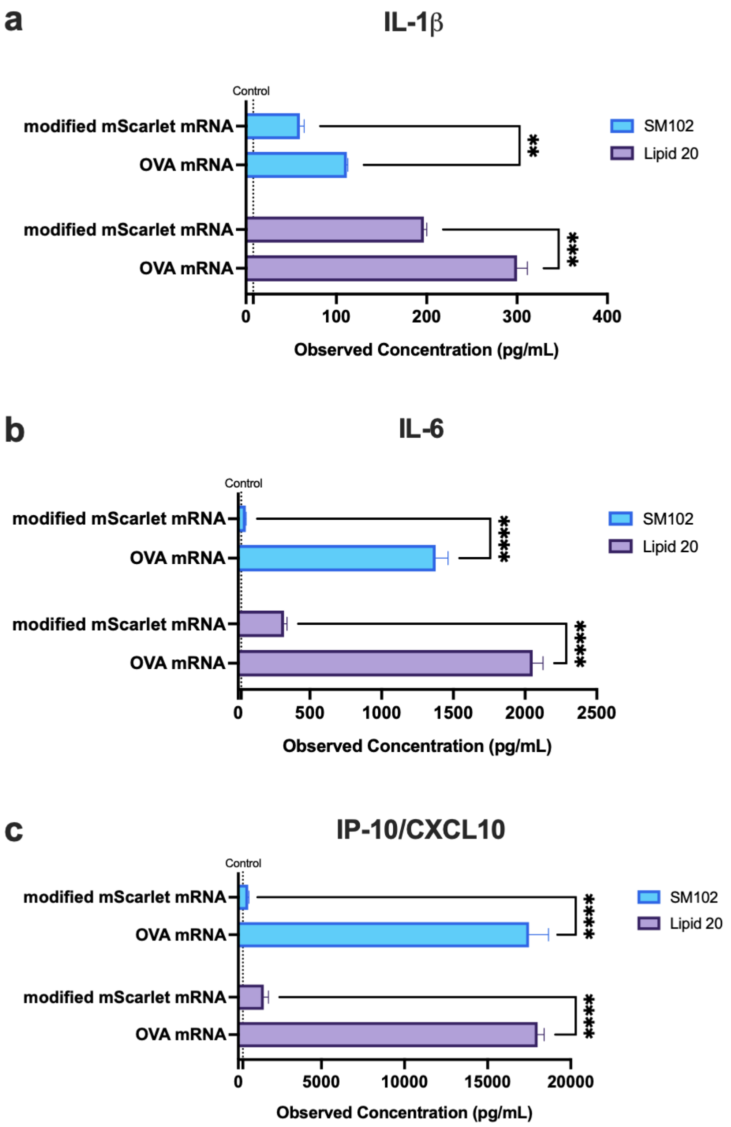


**Supplementary Figure 10. Cytokine responses in iMonocytes treated with unmodified OVA mRNA-LNPs.**

Cytokine secretion profiles of IL-1β **(a)**, IL-6 **(b)**, and IP-10/CXCL10 **(c)** following 72 hr-treatment of different mRNA-LNPs. Results were compared with those induced by the modified mScarlet mRNA-LNPs, and represent mean ± SEM, n = 3. *p < 0.05, **p < 0.01, ***p < 0.001, ****p < 0.0001 as analyzed by two-way ANOVA followed by the Tukey's multiple comparisons.
